# In-brain construction of receptor-based protease sensors by coupling ligand-directed chemistry and click chemistry

**DOI:** 10.1101/2024.05.23.594618

**Authors:** Seiji Sakamoto, Kazuki Shiraiwa, Mengchu Wang, Mamoru Ishikawa, Hiroshi Nonaka, Itaru Hamachi

**Author notes:** Correspondence should be addressed to H.N. and I.H. These authors contributed equally to this work (Seiji Sakamoto, Kazuki Shiraiwa).

## Abstract

The chemical modification of natural proteins in living systems is highly desirable toward the cutting-edge research in chemistry-biology interface. Recent advances in bioorthogonal protein modification have enabled the production of chemically functional proteins in cultured cell systems. However, few methods are applicable *in vivo* because of the complexity of the three-dimensional constructs of living systems with diverse, heterogeneous cell populations and flow systems filled with tissue fluids. Here, we report a genetic engineering-free method to modify receptor proteins with various probes in the living mouse brain by combining in-brain ligand-directed chemistry with bioorthogonal click chemistry, and propose a chemical guideline for the reaction design. The rapid and selective tethering of a set of fluorescent peptides to AMPA-type glutamate receptors (AMPARs) allowed the construction of receptor-based fluorescent sensors. These probes enabled mapping of the activity of matrix metalloproteinase-9 proximal to AMPARs in the living brain to be realized with high spatial resolution. Our strategy provides new opportunities for the precise analysis of particular *in vivo* microenvironments that has not been able to be addressed by conventional methods. Such analysis should contribute to the understanding of the molecular basis for complicated *in vivo* events, such as the regulation of neuroplasticity, the most important challenge in neuroscience.

## Introduction

Protein chemical modification is important in basic science and applied research.^1–8^ Various probes chemically tethered to target proteins allow the natural functions of the proteins to be expanded for use as bioimaging tools, biosensors, and bio-pharmaceuticals. Although the chemical modification of proteins has been often performed using purified proteins *in vitro*, precise modification in more natural environments, such as live cells and *in vivo*, is highly desirable for state-of-the-art research in chemistry-biology interface. Compared with *in vitro* chemistry, achieving protein selectivity and acceptable modification yields remain challenging because live cells and in living systems are spatially heterogeneous and contain multiple biomolecules under crowding conditions. Recent progress in bioorthogonal protein chemistry, including the incorporation of nonnatural amino acids by genetic code expansion coupled with a click reaction, and chemogenetic approaches using enzymes (such as Halo/SNAP)-tag based protein engineering, has enabled the production of chemically modified proteins in cultured cell systems.^9,10^ While these methods are powerful, to date very few methods are applicable to *in vivo* systems, which are three-dimensional (3D) constructs with diverse cell populations and a flow system of tissue fluids that are considerably unlike the two-dimensional (2D) closed systems of cells cultured on a petri dish.^11,12^

The brain has the most complex 3D structure of all *in vivo* systems. In the brain, extensive networks of neurons are tightly supported by glial cells, blood vessels, cerebrospinal fluid (CSF), and other substances to maintain brain homeostasis in both structure and function. The central neural system contains many synapses of 1 µm or less in size. In these synapses, neurotransmitter receptors play crucial roles in inter-neuron communication and the resultant sophisticated brain functions. Neurotransmitter receptors, such as glutamate receptors, are surrounded by other proteins and enzymes, such as other receptors, synaptic organizers, and proteases and peptidases, by which their functions are tightly regulated. Although such dynamically modulated molecular networks centered on the receptors are known to be involved in brain functions and disorders, the details have not been well deciphered because of the lack of analytical tools applicable to live brain systems. Chemistry-based methods are anticipated to uniquely contribute to the understanding of neural networks at the molecular level, but such approaches are still only in their infancy. We have recently reported in-brain ligand-directed acylimidazole (LDAI) chemistry as a method for the chemical modification of neurotransmitter receptors in the complex context of the brain.^13,14^ This method can be used to chemically label and visualize target receptors with high selectivity in the living mouse without genetic manipulation. While promising, there are still many challenges with this method, including the slow modification kinetics, the requirement for the time-consuming individual design and synthesis of a labeling reagent for each probe,^13–16^ and the lack of guidelines for the rational design of affinity-driven labeling reactions in the live brain. To address these issues, we sought to combine in-brain LDAI chemistry with bioorthogonal click reactions.^17,18^ This approach is designed to separate the anchoring step of a bioorthogonal reaction handle to the target receptor and the step for the modification of a functional probe, to expand the flexibility for probe selection. Because of the very limited examples of click reactions in the brain^19^ and the poor experimental data for the in-brain chemical modification of proteins, we investigated a series of LDAI labeling reagents and click reagents with varied physicochemical properties. Herein, we describe a design guideline, for both the chemical labeling (anchoring) of target receptors with a click handle in the brain and the modification of receptors with functional probes using click reagents. Using this guideline, we achieved the rapid functional modification of a target receptor protein in a living mouse brain without the use of genetic techniques, termed “chemical knock-in (KI)”. The chemical KI of fluorescent peptide probes to AMPARs allowed the construction of receptor-based fluorescent sensors in a live brain. These sensors enabled the successful mapping of the activity of matrix metalloproteinase-9 (MMP9) (a representative protease involved in the regulation of brain function) that was proximal to AMPARs, in the context of the 3D live brain.

**Figure 1.**
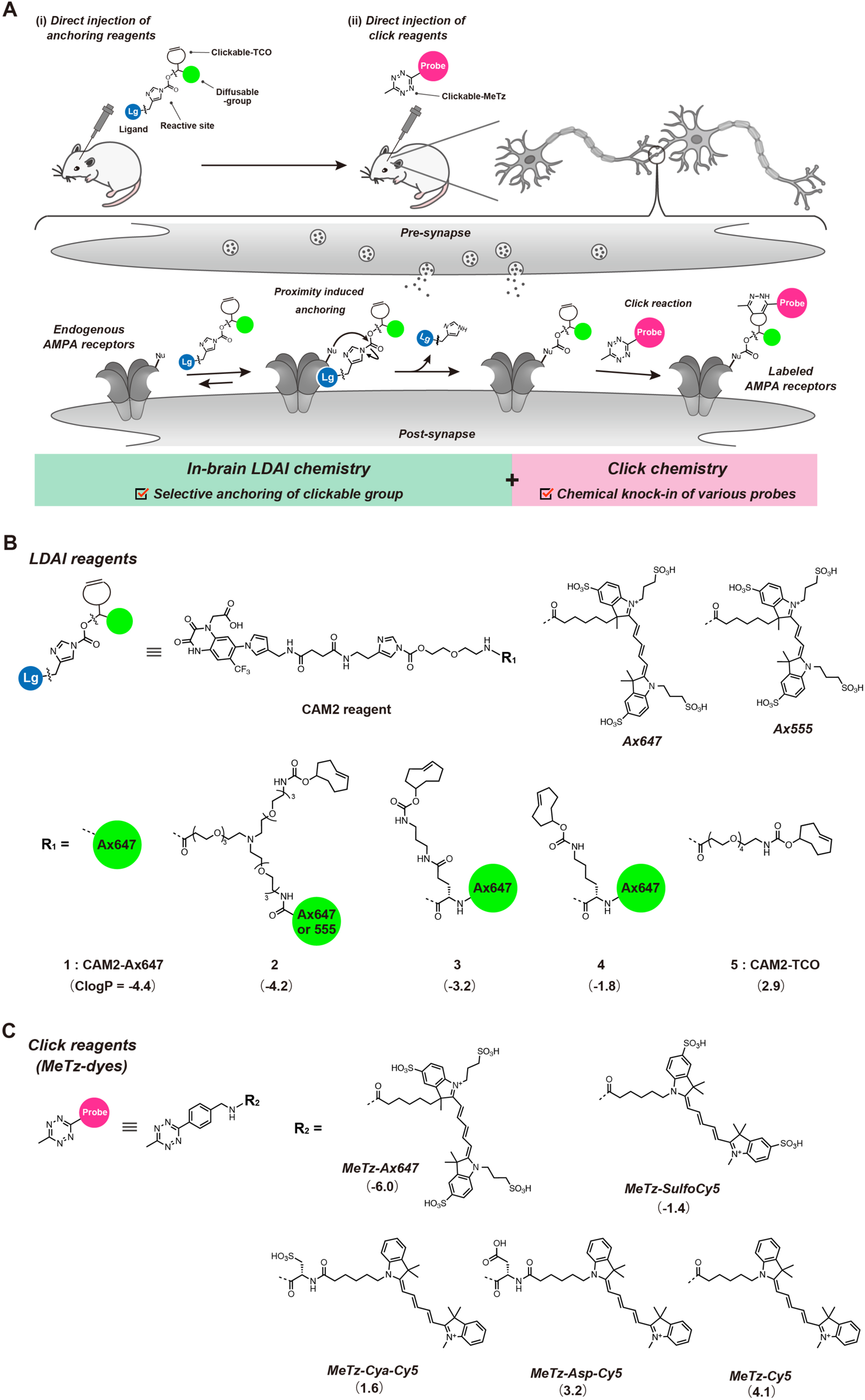
Chemical knock-in of functional probes to AMPA receptors in the living mouse brain. **A**, Schematic illustration of the chemical knock-in strategy used in the study. Nu, nucleophilic amino-acid residue. Lg, selective ligand for target receptor. TCO, *trans*-cyclooctene. MeTz, methyltetrazine. **B**, Chemical structures of LDAI reagents for anchoring TCO to AMPARs. The structures of Alexa Fluor 647 (Ax647) and Alexa Fluor 555 (Ax555) shown are the putative structures. **C**, Chemical structures of MeTz-based labeling reagents for the click reaction with a TCO group attached to AMPAR.

## Results

### Chemical KI strategy using in-brain LDAI and click chemistry

In established bioorthogonal protein modification using click chemistry, the incorporation of a nonnatural reaction (clickable) handle, such as an azide, is required for a protein-selective reaction. A state-of-the-art genetic expansion protocol using a suppressor codon can enable the incorporation of such a handle into a target protein in live cell systems.^20–22^ However, the robust application of this strategy in live animals, such as mice, is still very difficult and applications in the brain have been limited to date. While the metabolic incorporation of a clickable substrate is an alternative method, this approach generally suffers from insufficient selectivity toward the target protein.^23–25^

In our chemical KI approach, we used our LDAI chemistry to incorporate a clickable handle into a target receptor in the living brain. This anchoring step was followed by click chemistry to realize the functional modification of a receptor protein (labeling step). The approach is summarized in Figure 1A. Because LDAI chemistry can modify endogenous receptors, the chemical KI does not require any genetic manipulation. Chemical KI should be particularly beneficial for targeting structurally complicated oligomeric proteins in which the subunit balance is often disrupted by genetic engineering. In addition, genetic engineering techniques are still not well established in many animals. We here targeted the AMPARs, a major neurotransmitter receptor involved in fast excitatory neurotransmission, for a proof-of-principle study of chemical KI in the brain. As the click handle, we selected *trans*-cyclooctene (TCO), which reacts with many tetrazine (Tz) derivatives with fast kinetics (1–10^6^ M^-1^s^-1^ *in vitro*)^26–28^, to enable the rapid introduction of functional molecules to AMPARs *in vivo*. If these two steps can be successfully performed in the animal brain, the selective functionalization of endogenous AMPARs, that is the chemical KI of designer probes for a target protein, will be realized.

However, the conditions for the chemical reactions in the live brain (*in vivo*) are greatly different from those in live cells cultured in 2D Petri dishes (*in vitro*). In addition to the complexity of the 3D-structure with diverse and heterogeneous cell populations, the brain also has a flow system with constant irrigation and drainage of CSF.^29^ Under such conditions, chemical reagents are gradually removed from the brain, which can hinder the distribution of the reagents over a wide area of the brain. In our recent studies of in-brain LDAI chemistry, we found that the method of administration was critical for efficient delivery of the LDAI reagents to the whole brain, and the direct injection of the reagent into the lateral ventricle (LV) gave excellent results. After injection into the LV, the labeling reagent was diluted by CSF constantly produced from the choroid plexus of the LV, and the CSF flow from the LV to various areas allowed the labeling reagent to diffuse to tissues throughout the whole brain. Because the parameters required for efficient chemical reactions in the brain are expected to be different from those *in vitro*, we sought to compare the chemical reactivity between the conventional live cell systems and the live brain, using identical sets of LDAI and click reagents. Furthermore, considering that there are no design guidelines for either the anchoring step using LDAI or the protein modification (or labeling) step by click reaction in the live brain, we examined the reaction profiles of these two steps separately in detail.

### Selective anchoring of TCO groups to target proteins by ligand-directed chemistry

For anchoring the clickable TCO moiety to AMPARs, different LDAI reagents for AMPARs (chemical AMPAR modification reagents: CAM2) were designed and synthesized, based on the molecular structure of CAM2-Ax647 (**1**), which has previously been shown to have an excellent labeling capability in live mouse brains.^13^ The three labeling reagents (**2**, **3**, and **4**) contained Ax647 and TCO that were connected with different spacer units, an *N*-branched structure, Glu, or Lys, respectively (Figure 1B). We also employed CAM2-TCO (**5**) lacking Ax647 and CAM2-Ax647 (**1**) lacking TCO as control compounds.^30,31^ These five compounds had distinct physicochemical properties, which could be indexed by the values for the calculated logP (ClogP), a parameter of hydrophobicity, which were **1** (-4.4), **2** (-4.2), **3** (-3.2), **4** (-1.8), and **5** (2.9).

We first evaluated the selectivity and efficiency of the TCO-anchoring reaction in HEK293T cells expressing GluA2 (a subunit of the AMPA receptor) (Figure 2A and 2B). A CAM2 reagent (1 µM) was added to live HEK293T cells at 37°C, followed by incubation for 3 h and cell lysis. The lysates were analyzed by SDS-PAGE in-gel fluorescence. In the case of CAM2-TCO (**5**), the lysate was mixed with **Tz-Ax647** prior to SDS-PAGE analysis, as previously described.^30^ As shown in Figure 2B, all the CAM2 reagents gave a single band corresponding to GluA2, indicating the high AMPAR selectivity in the TCO anchoring step. In addition, the intensities of fluorescence bands derived from the labeled Alexa647 were almost comparable to each other, indicating that the anchoring efficiency was not dramatically different between these CAM2 reagents (**1–5**), within a 30% variation.

The selectivity and efficiency of the TCO anchoring step using these reagents were also evaluated in living mouse brains (Figure 2A and 2C). Mice were injected with 4.5 µL of each labeling reagent (100 µM) into the LV and 24 h after injection, the cerebellum, which is relatively distant from the LV and rich in endogenous AMPA receptors, was isolated. The homogenates were subjected to SDS-PAGE analysis. The homogenates of the mouse brains injected with CAM2-TCO (**5**) were treated with **MeTz-Ax647** prior to SDS-PAGE. A single band corresponding to a subunit of the AMPAR was detected for the CAM2 reagents (**1–4**) in the in-gel fluorescence analysis, indicating high AMPAR selectivity. In contrast, negligible bands were observed for CAM2-TCO (**5**), which suggested that the anchoring reaction had scarcely occurred. The anchoring efficiency was compared from the band intensities, CAM2-Ax647 (**1**) lacking TCO had the highest intensity, and for the TCO-bearing CAM2 derivatives, the efficiency declined gradually from the derivatives with an *N*-branch (**2**) to Glu (**3**), Lys (**4**), and CAM2-TCO (**5**). Figure 2D shows a plot of the anchoring efficiency against the ClogP values of the five CAM2 reagents. The efficiency of the reaction in the live brain corresponded to the ClogP value of the CAM2 reagent; that is, compounds with more negative ClogP values resulted in more efficient TCO anchoring to the AMPARs. In sharp contrast, the anchoring efficiency in HEK 293T systems (*in vitro*) was not strongly related to the ClogP values (Figure 2C and 2D). For example, CAM2-TCO (**5**) failed to tether the TCO group with the AMPA receptors in the brain, whereas this reaction was successful *in vitro*.

The spatial distribution of the anchoring reaction was subsequently examined using a confocal Laser Scanning Microscope (CLSM) of the brain sections. In brain slices injected with the CAM2 reagent **2**, fluorescence from Ax647 was observed from the hippocampus, cerebellum, cortex, and striatum, where endogenous AMPARs are highly expressed (Figure 2E). Although the cerebellum was distant from the administration site of the reagents, a fluorescent signal was clearly observed, revealing the excellent in-brain diffusivity and permeability of **2**. Importantly, the fluorescence derived from the anchored Ax647 showed a pattern similar to that found with co-immunostaining using anti-GluA2 antibodies (Figure 2E) and the high-resolution images of the cortex area showed numerous bright spots of approximately 1 µm in size. These bright spots also overlapped well with the anti-GluA2 fluorescence, implying that these spots indicated neuronal spines where endogenous AMPARs were accumulated (Figure 2F).

Considering that the LDAI reagents need to reach target AMPARs in several distinct regions of the brain, the diffusion and permeability of the CAM2 reagents through heterogenous brain tissues could be crucial factors for efficient ligand-directed TCO anchoring in the live brain. These factors may be partly expressed by the ClogP value and this value was found to be a helpful indicator in the design of chemical reagents for in-brain ligand-directed receptor labeling. Compared with the live brain, the cultured cell system was rather simple and thus the anchoring step might be less sensitive to the parameters reflected by the ClogP value in this system.

**Figure 2.**
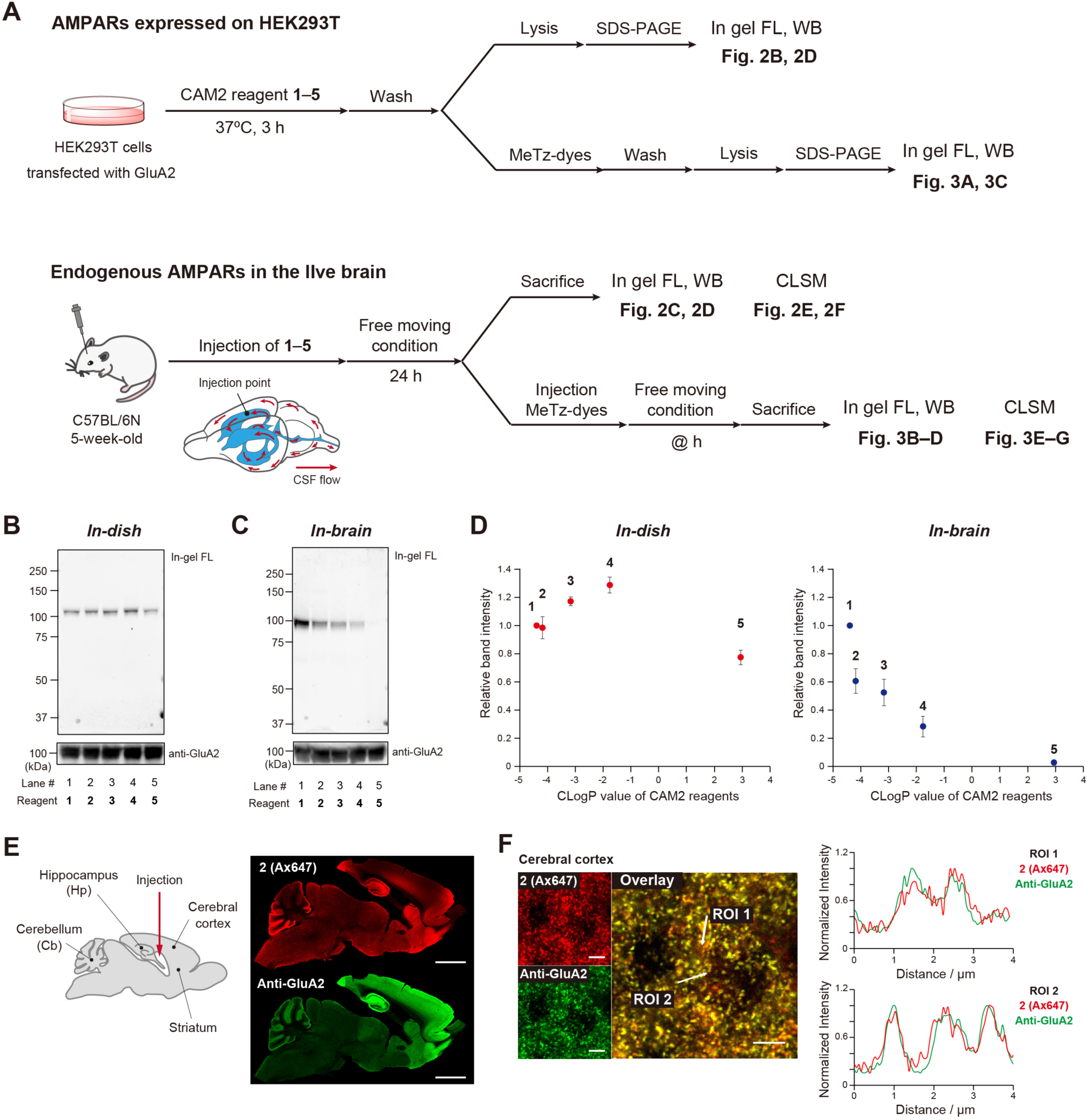
TCO anchoring to AMPA receptors using LDAI chemistry. **A,** Experimental workflow for the results shown in Figure 2 and Figure 3. **B**, In-gel fluorescence and western blotting (WB) analysis of the labeling reaction with AMPARs expressed with GluA2 under in-dish conditions. **C**, In-gel fluorescence and WB analysis of the labeling reaction with endogenous AMPARs in the live brain. **D**, Comparison of the relative labeling efficiency between in-dish and in-brain conditions. Data are presented as means ± SEM. **E**, Fluorescence images of labeled brains with **2 (Ax647)** and colocalization analysis with anti-GluA2. A mouse was transcardially perfused with a solution of 9% glyoxal/8% acetic acid (pH 4.0) 24 h after LV injection of **2**. The brain was isolated and sectioned by cryostat (50 µm thick sections). The sections were immunostained with anti-GluA2. Imaging was performed using a CLSM equipped with a 10× objective. Scale bar: 2 mm. **F**, High-resolution confocal image of the cerebral cortex with 100× lens. Scale bar: 5 µm.

### Highly selective and fast chemical KI using click chemistry

We next investigated the click reaction (probe labeling) step. According to previous reports of the fast reactivity with the TCO moiety and good stability *in vivo*, methyltetrazine (MeTz) was selected as the clickable moiety and a set of fluorescent (Cy5 core) derivatives with various ClogP values was prepared as follows: **MeTz-Ax647** (ClogP = –6.0), **MeTz-SulfoCy5** (–1.4), **MeTz-Cya-Cy5** (1.6), **MeTz-Asp-Cy5** (3.2), and **MeTz-Cy5** (4.1) (Figure 1B). For the TCO anchoring to AMPARs, CAM2 (**2)** bearing Ax555 instead of Ax647 was used in both live cells and the live brain study to enable the use of two distinct wavelengths to evaluate the click reaction and the TCO anchoring separately.

We conducted the first anchoring step of AMPA receptors transiently expressed in HEK293T cells according to the protocol shown in Figure 2A, [**2 (Ax555)**, 1 µM, 37°C, 3 h]. Subsequently, the cells were incubated with each of the five MeTz derivatives (1 µM, 1 h), followed by washing and cell lysis and then the lysates were subjected to in-gel fluorescence analysis. The reaction of MeTz with the TCO-tethered AMPARs could be monitored by the Cy5 emission. As shown in Figure 3A, single bands exhibiting almost identical emission intensity were detected for **MeTz-Ax647**, **MeTz-SulfoCy5**, **MeTz-Cya-Cy5**, and **MeTz-Asp-Cy5**. In the case of **MeTz-Cy5**, the nonspecific labeling of proteins other than AMPA receptors was also observed. The observed intensity indicated that the click reaction in HEK cells was not dependent on the ClogP value of the MeTz derivatives, which displayed a similar trend to that for the LDAI-based TCO anchoring step.

In the live mouse brain system, the anchoring step and the click reaction step that followed were performed according to the protocol shown in Figure 2A. The TCO-anchoring reagent **2 (Ax555)** (100 µM, 4.5 µL) was injected into the LV of mice under anesthesia and after 24 h, a MeTz probe was injected into the LV (100 µM, 4.5 µL). The cerebellum was isolated after 24 h and the homogenates were subjected to SDS-PAGE in-gel fluorescence analysis. **MeTz-Ax647** (ClogP = –6.0) and **MeTz-SulfoCy5** (–1.4) showed greater click reaction efficiency, relative to the other compounds, and almost no bands were observed for **MeTz-Cy5** (Figure 3B). Figure 3C shows that MeTz derivatives having a negative ClogP value resulted in better modification yields than those having a positive ClogP value. Compared with the in-brain LDAI chemistry, the click reaction step was less sensitive and rather more tolerant to compounds with varying ClogP values. Overall, the reaction efficiency in the brain system was largely correlated with the ClogP value of the reagents in both the anchoring and modification steps, whereas the efficiency in the in-dish cultured cell system did not show any appreciable correlation with the ClogP values.

Using the efficient in-brain chemical KI tools [**2 (Ax555)** and **MeTz-Ax647**], we next determined the modification kinetics of endogenous AMPARs in the live brain. Mice bearing TCO-tethered AMPA receptors were injected with **MeTz-Ax647** (100 µM, 4.5 µL) into the LV. The cerebellum was isolated at various time points after the injection and the reaction efficiency was assessed by the in-gel fluorescence band intensity (Figure 3D). It is clear from Figure 3D that the click reaction reached saturation in less than 60 min, indicating that the high reactivity of the inverse electron-demand Diels–Alder (IEDDA) reaction between TCO and MeTz resulted in a rapid reaction in the live mouse brain, which enabled chemical KI of the fluorescent probe Ax647 into the target AMPA receptors within 1 h. We also evaluated the duration of **MeTz-Ax647** in the mouse brain without TCO-anchored AMPARs. The half-life of **MeTz-Ax647** was ca. 0.5 h (Supplementary Figure 1), clearly indicating that the live brain had a flow system that enabled the chemical reagents to be removed gradually. We believe that consideration of this flow system is critically important in the design of in-brain chemical reactions with high efficiency.

Finally, we conducted imaging analysis of the brain after the click (modification) reaction (Figure 3E–3G). For **MeTz-Cy5**, strong fluorescence was observed in very limited areas close to the LV, the **MeTz-Cy5** injection site, whereas very weak fluorescence was observed in the hippocampus and cerebellum regions where AMPARs are highly expressed (Figure 3E). Combined with the in-gel fluorescence results, the strong signal close to the LV areas was believed to mainly result from the nonspecific binding or adsorption of **MeTz-Cy5**. For **MeTz-Ax647** and **MeTz-SulfoCy5**, a broad distribution of fluorescence signals was observed throughout the brain areas expressing endogenous AMPARs (Figure 3E and Supplementary Figure 2). Imaging at a higher magnification clearly revealed that the fluorescent bright spots, of 1 µm or less in size, of TCO-anchored AMPARs (Ax555) in the spines were well merged with the signals derived from the click modification (Ax647 emission, Figure 3F). Interestingly, these labeled signals were substantially retained even after clearing of the whole brain by the CUBIC method^32^ and these punctate signals could be observed in 3D mode without 2D-section preparation (Figure 3G). These imaging analyses demonstrated that the diffusivity and permeability of the MeTz derivatives had a critical impact on the in-brain click reaction, which was reflected in the ClogP values, as was the case for the in-brain LDAI chemistry.

**Figure 3.**
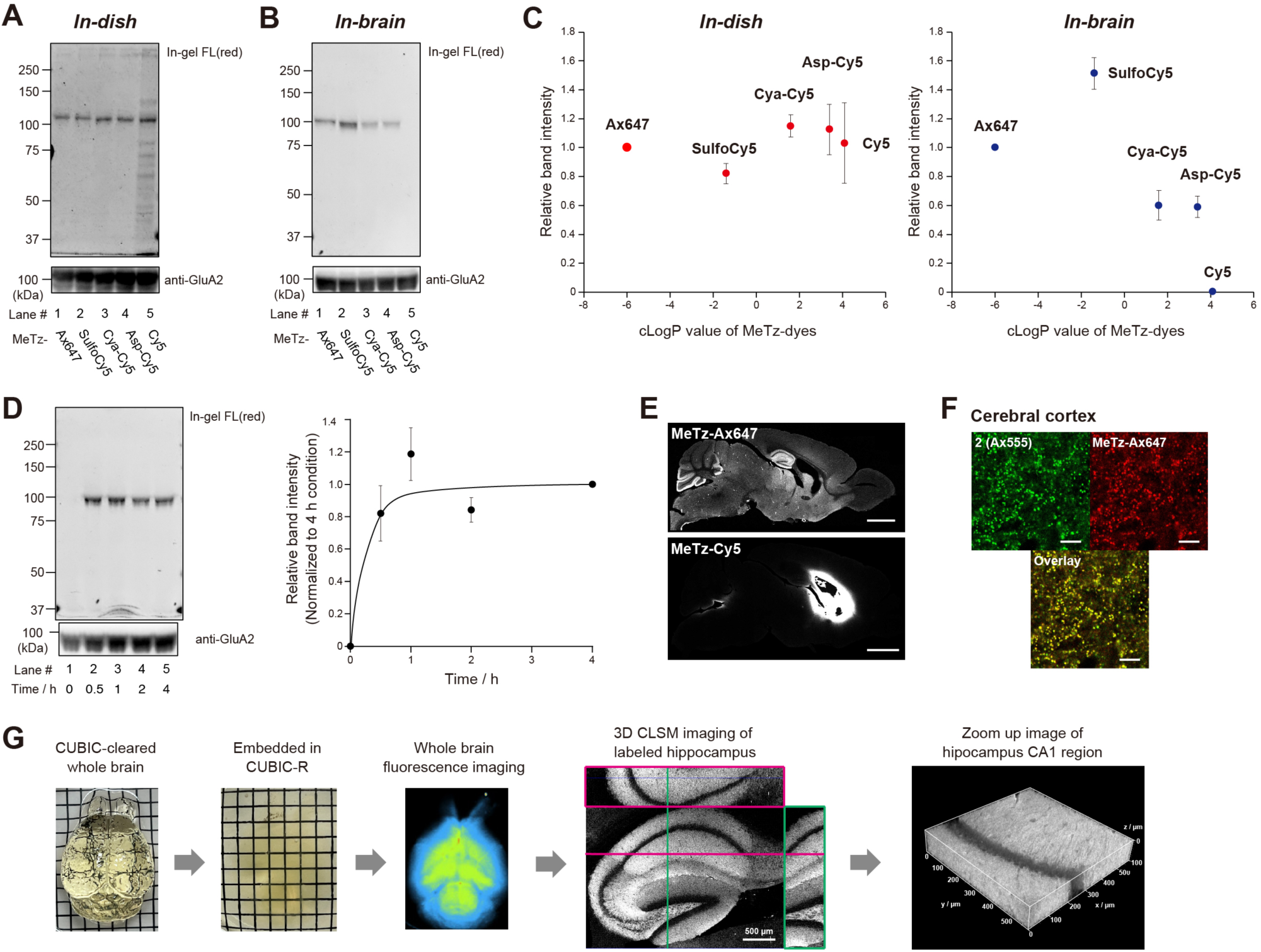
Probe modification of AMPA receptors by click chemistry. To facilitate the analysis of the click reaction, the TCO anchoring reagent **2 (Ax555)** was used in the experiments shown in Figure 3 and thereafter. **A**, In-gel fluorescence and WB analysis of the click reaction with AMPARs expressed with GluA2 in the in-dish conditions. **B**, In-gel fluorescence and WB analysis of the click reaction with endogenous AMPARs in the brain. **C**, Comparison of the labeling efficiency between the in-dish and in-brain conditions relative to **MeTz-Ax647**. Data are presented as means ± SEM. **D**, Kinetic analysis of in-brain click reaction with TCO-anchored AMPA receptors. Relative labeling efficiency normalized by the in-gel fluorescence intensity after 4 h of click reaction. Data are presented as means ± SEM. **E**, Fluorescence images of sagittal brain sections labeled by the click reaction with **MeTz-Ax647** or **MeTz-Cy5**. A mouse was additionally injected with **MeTz-Ax647** or **MeTz-Cy5** (100 µM, 4.5 µL), 24 h after LV injection of **2 (Ax555)** (100 µM, 4.5 µL). Then, 24 h after the LV injection of the MeTz derivatives, the mouse was transcardially perfused with a 4% PFA/PBS(–) solution. The brain was isolated and sectioned by cryostat (50 µm thick sections). Imaging was performed using a CLSM equipped with a 10× objective. Scale bar: 2 mm. **F**, Zoomed image of the cerebral cortex shown in Figure 3E. Imaging was performed using a CLSM equipped with a 100× objective. Scale bar: 5 µm. **G**, Fluorescence imaging of a tissue-cleared brain labeled with **2 (Ax555)** (100 µM, 4.5 µL; left and right LV) and **MeTz-Ax647** (100 µM, 4.5 µL; left and right LV). Fluorescence imaging of CUBIC-cleared whole brain was performed using FUSION, a chemiluminescence/fluorescence imaging system (excitation wavelength 600–650 nm, detection wavelength 690–720 nm). Three-dimensional imaging of the hippocampus area was performed using a CLSM equipped with a 10× and a 20× objective. FRET signals beween Ax647 (click labeling) and Ax555 (TCO anchoring) were observed with the laser excitation at 561 nm and emission at 655–770 nm.

### Construction of an AMPAR-based fluorescent MMP9 sensor in the live brain and evaluation of MMP9 activity proximal to the receptors

Having an efficient in-brain chemical KI method in hand, we applied this method to the construction of receptor-based protease sensors. In the brain, proteases such as MMP9 play crucial roles in the regulation of synaptic function by reorganizing the extracellular matrix, degrading synaptic organizers, and activating cytokines.^33,34^ However, the molecular details of the activity of particular proteases are not fully understood, mainly because there are no valid methods capable of directly analyzing protease activity with high spatial resolution in a live brain. Fluorescent protease biosensors that can monitor the protease activity *in situ* are expected to be useful for such analysis, and thus we constructed AMPAR-based sensors via chemical KI of the corresponding protease substrates (Figure 4A). In detail, AMPARs anchored with Ax555 and TCO were modified with a variety of Ax647-appended peptides (protease substrates) via the in-brain click reaction. When a protease cleaves the peptide tethered to the AMPARs, the Ax647 emission will decrease, resulting in an increase in the ratio of Ax555/Ax647. Given that the cleavage site is < 10 nm from the surface of the AMPAR, this sensor is able to detect protease activity that is proximal to the receptor. The fluorescent peptide substrates for MMP9 (MMP9 probes) were designed based on previous reports, as shown in Figure 4B.^35,36^ MeTz and the fluorescent Ax647 were attached at the *C*-, and *N*-termini of the peptides, respectively. Two different sequences with high and low reactivity for MMP9 were prepared (**MMP9-Leu**, **MMP9-Abu**), together with **MMP9-DLeu** where the L-Leu of the cleavage site was replaced with the D-isomer, as a non-cleavable control. Furthermore, an Ax647-appended peptide substrate for cathepsin B (CatB), which is an intracellular protease mainly localized in endosomes, was designed in the same manner (CatB probe, Figure 4B).^37^ We separately confirmed the enzymatic cleavage of these probes by HPLC analysis (Supplementary Figures 3 and 4).

First, we prepared a fluorescent MMP9 sensor for the AMPAR scaffold transiently expressed in HEK293T cells and investigated whether the sensor could detect MMP9 protease activity. Live cell CLSM imaging showed that Ax647 fluorescence derived from the **MMP9-Leu** peptide was observed on the cell surface, together with Ax555 emission (Figure 4C, 0 min), indicating that the AMPARs successfully underwent modification with the MMP9-peptide by click chemistry. When a solution of collagenase containing MMP9 (gelatinase B, type IV collagenase) was added to the HEK293T cells, the emission intensity of Ax647 decreased without a change in the Ax555 emission (Figure 4C, 30 and 60 min). Such fluorescence changes did not occur with **MMP9-DLeu** modification (Figure 4D). Ratiometric analysis of the fluorescence intensity of Ax555/Ax647 at the cell surface allowed the evaluation of the MMP9 activity more quantitatively (Figure 4E). These results indicated that an AMPAR-based MMP9 fluorescent sensor was successfully constructed by chemical KI in a cultured cell system.

**Figure 4.**
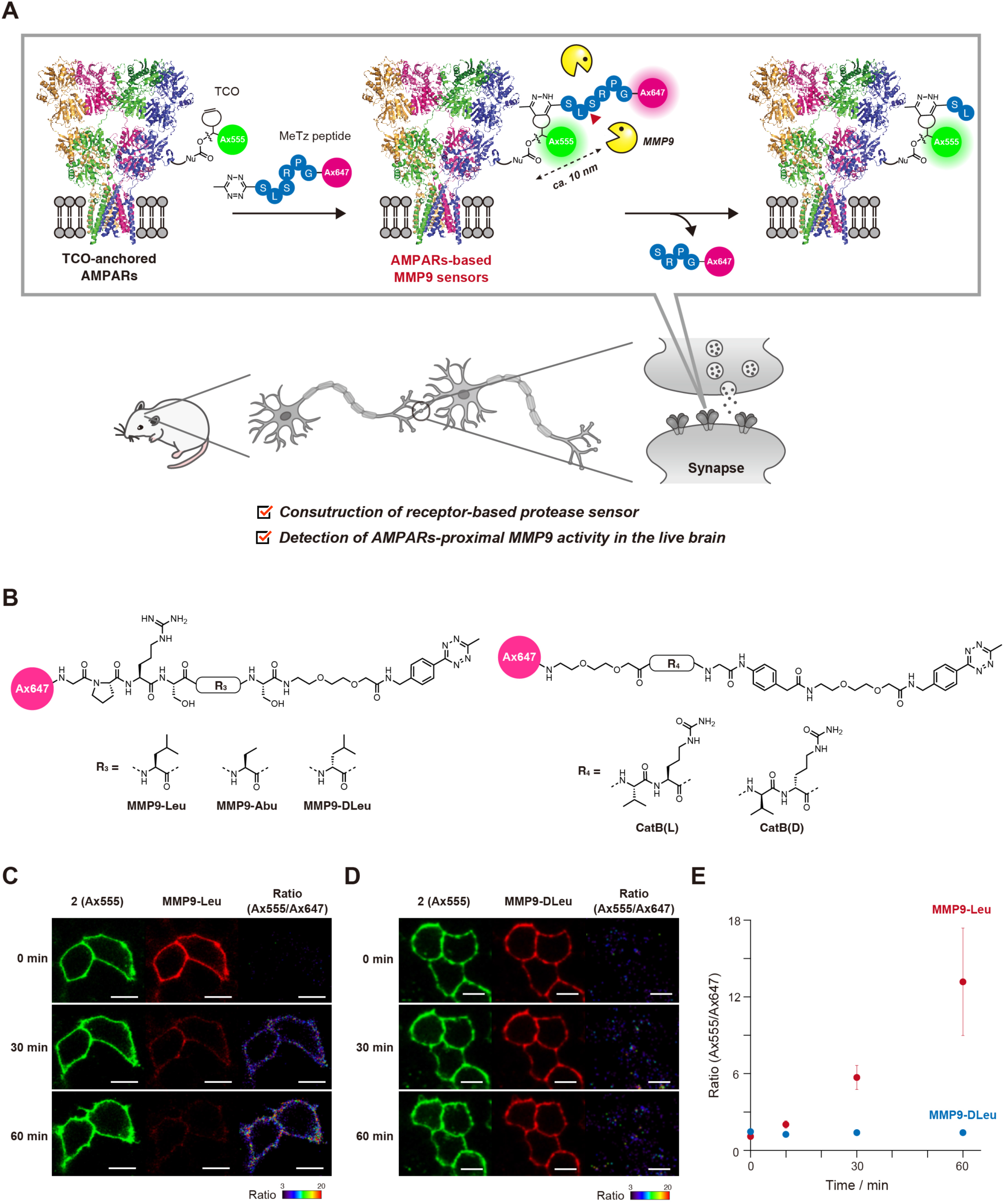
Construction and evaluation of an AMPAR-based fluorescent MMP9 sensor in HEK293T cells. **A**, Schematic illustration of the analysis of MMP9 activity proximal to AMPARs. **B**, Chemical structure of fluorescent peptide probes for the detection of MMP9 and CatB activity. **C–D**, Time course analysis of MMP9 activity using an AMPAR-based fluorescent MMP9 sensor. CLSM observations were conducted at different times after the addition of collagenase. Fluorescence imaging was performed using a Leica CLSM equipped with a 40× objective. Scale bar: 10 µm. **E**, The plot of Ax555/Ax647 values on the cell surface versus time. Region of interest (ROI) (n = 5). Data are presented as means ± SEM.

Finally, we attempted to construct a fluorescent MMP9-sensor in a live brain. After anchoring AMPA receptors with TCO and Ax555 by LV injection of **2** (Ax555) in a mouse brain, an Ax647-appended peptide (MMP9 or CatB probe) was subsequently injected to the LV site. The hippocampus was isolated 6 h after the second injection, and the homogenates were analyzed by SDS-PAGE in-gel fluorescence. A single Ax647-fluorescence band was observed at ca. 100 kDa corresponding to the endogenous AMPAR subunit, indicating the selective modification of endogenous AMPARs with these peptide probes (Figure 5A). The hippocampal samples labeled with **MMP9-Leu** or **MMP9-Abu** showed weak fluorescence bands (Lanes 1 and 4), relative to the band of the sample labeled with **MMP9-DLeu** (Lane 3). Importantly, the fluorescence band of the sample after co-injection with **MMP9-Leu** and ilomastat, an MMP inhibitor, retained intensity that was almost comparable to that of the **MMP9-DLeu** sample (Lane 2). These results were in good agreement with the results for the cleavage reactivity of the MMP9 probes *in vitro* using hippocampal homogenates (Supplementary Figure 5). It was concluded that the relative intensity of the in-gel fluorescence band reflected the MMP activity; that is, the decrease in the fluorescent bands for **MMP9-Leu** and **MMP9-Abu** indicated high MMP9 activity in the hippocampal region. In contrast, the differences in the signals between the samples labeled with the **CatB(L)** and **CatB(D)** probes were small (Figure 5A, Lanes 5 and 6), which suggested that the CatB activity was low in the extracellular region of the hippocampus.

The fluorescence images of cryosections were expected to provide spatial information on the protease activity in the whole brain, and we observed fluorescence in sagittal brain sections, 6 h after injection of **MMP9-Leu** using a 5× objective lens. Figure 5B shows that there was a negligible signal from Ax647 in the hippocampus and substantial fluorescence in the cerebellum, relative to the signals of the corresponding regions after **MMP9-DLeu** injection, suggesting distinct activity of MMP9 depending on the region. In these experimental setups, however, the decrease in the Ax647 signal may have been caused by two possible scenarios; that is, the peptide probe was cleaved before reaching the AMPARs and/or the probe was cleaved after tethering to the AMPARs. To explicitly evaluate the cleavage of peptides attached to the AMPARs, we performed snapshot CLSM imaging of brain slices after different incubation times. Perfusion fixation was conducted 3, 6, and 9 h after the probe injection. In the hippocampal regions modified with **MMP9-Leu**, almost no Ax647 fluorescence was observed even after incubation for 3 h (Figure 5C, left panels). The samples modified with **MMP9-Abu**, in contrast, showed definite Ax647 fluorescence at 3 h and a gradual decrease over incubation for 6 and 9 h, resulting in an increase in the Ax555/Ax647 ratio (Figure 5C, middle panels). In contrast, such changes were not observed for the samples injected with **MMP9-DLeu** (Figure 5C, right panels). These results clearly indicated that with **MMP9-Abu**, a substantial amount of the MMP9 probe was cleaved after being tethered to AMPARs. This result may reflect the lower reactivity of **MMP9-Abu** for MMP9 (compared with **MMP9-Leu**), and the slow kinetics allowed **MMP9-Abu** to reach AMPARs prior to the cleavage. This **MMP9-Abu** probe with moderate cleavage properties is more appropriate for the investigation of the MMP9 activity proximal to a receptor of interest and we used this probe in the following studies. Note that, as expected from the results shown in Figure 5A, there was little change in the fluorescence images of samples labeled with the **CatB(L)** probe and those labeled with the **CatB(D)** probe (Supplementary Figure 6).

The magnified CLSM images showed the MMP9 activity at a higher spatial resolution. In the hilus of the hippocampal dentate gyrus, the signals from Ax555 and Ax647 fluorescence derived from the AMPARs and the **MMP9-Abu** probes, respectively, were both observed as bright puncta of 1 µm or less in size and the images were well merged (Figure 5D), which indicated that the MMP9 probe was indeed tethered to AMPARs located in the dendritic spines (Supplementary Figure 7). Importantly, longer incubation times (from 3 to 9 h) decreased the Ax647 fluorescence and increased the value of the Ax555/Ax647 ratio in many of the spines in the samples labeled with **MMP9-Abu** (Figure 5D and 5E), strongly suggesting that MMP9 had penetrated into the synaptic cleft, closely approached the AMPARs (within 10 nm), and then cleaved the MMP9-substrate peptide. This result may be consistent with a previous report suggesting that MMP9 was responsible for the cleavage of synaptic organizers (such as neuroligins).^38^ To our knowledge, this is the first report of images that capture the MMP9 activity proximal to AMPARs in the hippocampal excitatory synapses of a live brain, which cannot be achieved using conventional methods. CLSM observations were also performed in the cerebellum where AMPARs are highly expressed. In the fluorescence imaging of the whole cerebellum, the Ax647 fluorescence derived from AMPAR-based MMP9-Abu sensors remained considerably high, 9 h after injection of the probe (Figure 5F). The high magnification images of the molecular layer region also showed that many of the Ax647 bright spots remained intact over 9 h of incubation (Figure 5G). When we compared the time-profile of the Ax555/Ax647 ratio between the hippocampus and cerebellum, at puncta-level resolution, the MMP9 activity proximal to AMPARs in the cerebellum was observed to be approximately 6-fold lower than that in the hippocampus area (Figure 5E and 5H). This is the first experimental evidence to showing MMP9 activity in the live brain is greatly dependent on the region.

**Figure 5.**
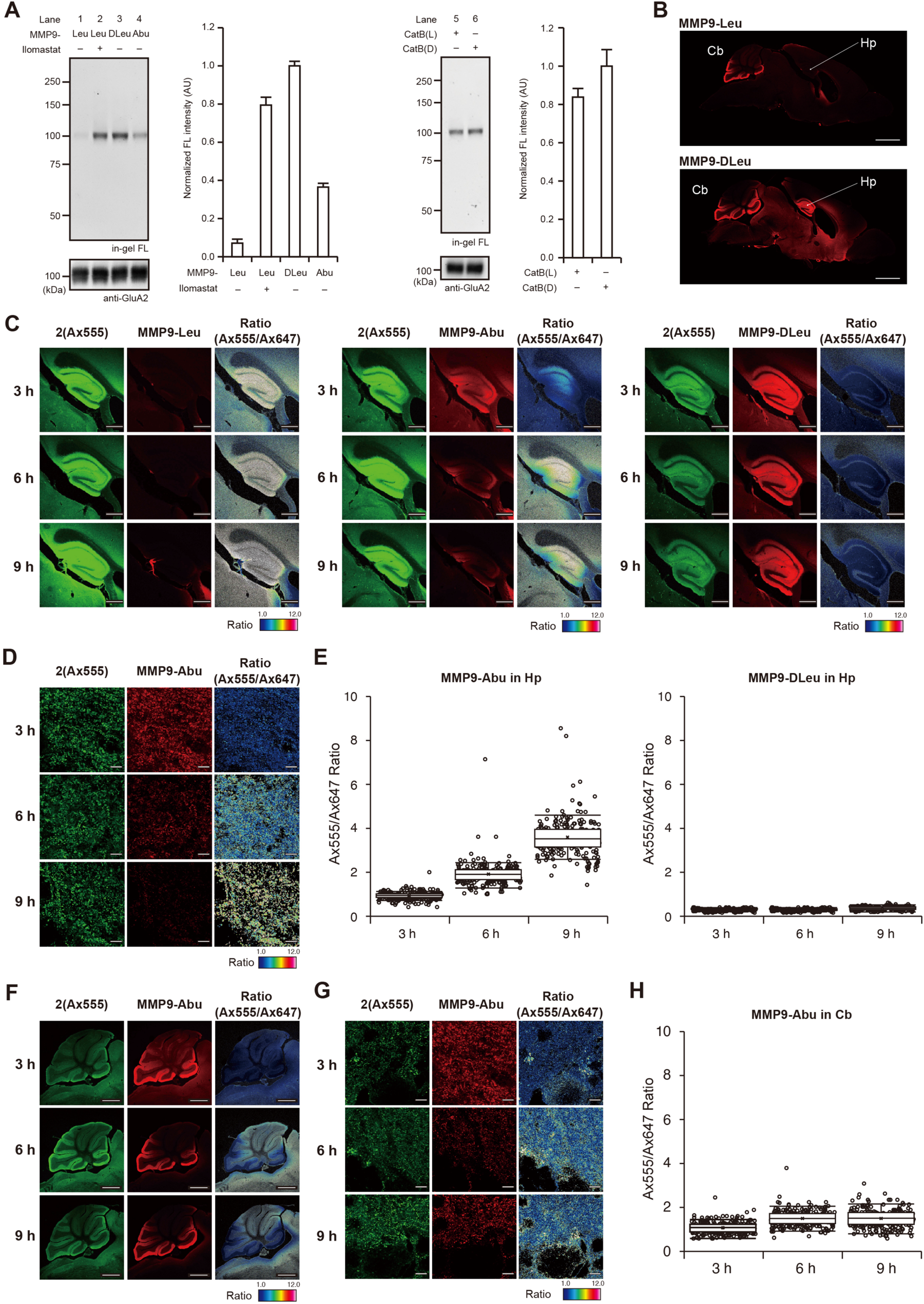
Construction and evaluation of an AMPAR-based fluorescent MMP9 sensor in a living mouse brain. **A**, Comparison of in-gel fluorescence band intensity at 6 h after the injection of peptide probes (100 µM, 4.2 µL). **B**, Fluorescence images of sagittal brain sections labeled by the click reaction with **MMP9-Leu** or **MMP9-DLeu**. A mouse was additionally injected with **MMP9-Leu** or **MMP9-DLeu** (100 µM, 4.2 µL), 12 h after LV injection of **2 (Ax555)** (100 µM, 4.2 µL). Then, 6 h after LV injection of MeTz derivatives, the mouse was transcardially perfused with a 4% PFA/PBS(–) solution. The mouse brain was isolated and sectioned by cryostat (50 µm thick sections). Imaging was performed using a CLSM equipped with a 5× objective (NA = 0.25). Scale bar: 2 mm. **C-F**, Detection of MMP9 activity by CLSM. **C**, Hippocampus area, 5× objective, scale bar: 0.5 mm. **D**, Hilus area of the hippocampal dentate gyrus, 63× objective (NA = 1.4), scale bar: 5 µm. **E**, Boxplot of the Ax555/Ax647 ratio values of the fluorescent bright spots in the hilus area of the hippocampal dentate gyrus. The number of ROIs was 240 for each of the images. **F**, Cerebellum area, 5× objective, scale bar: 1 mm. **G**, Area near Purkinje cells, 63× objective, scale bar: 5 µm. **H**, Boxplot of the Ax555/Ax647 ratio values of the fluorescent bright spots in the posterior lobe IX of the cerebellum. The number of ROIs was 240 for each of the images.

## Discussion

Over the decades, a variety of chemical methods for protein bioconjugation have been developed.^1–8^ However, the modification of a target protein in living animals has rarely been achieved, mainly because of the poor protein selectivity and inefficient reactions under the complex conditions present in living systems. In the present study, we have developed a strategy for the chemical KI of functional synthetic molecules into a target receptor in a live mouse brain with no genetic manipulation. The power of the chemical KI method was clearly demonstrated by the construction of AMPAR-based MMP9 protease sensors, which enabled, for the first time, the mapping of MMP9 activity in proximity (∼10 nm) to AMPARs in the postsynapses with high spatial resolution in a live brain context.

This entirely chemistry-based method was realized on the basis of our interesting finding of a design guideline for efficient chemical reactions in the live brain. In both steps of the method, employing LDAI and click reagents in the brain, we discovered an unprecedented correlation between the ClogP values of the reagents and the reaction efficiency that was not observed the reactions in cell culture systems. This result implied that the ClogP value may be a useful indicator for the design of chemical reactions in the brain (and probably elsewhere in *in vivo* systems). Our study also indicated that the diffusivity and the tissue-penetration properties of chemical reagents can be related to the target selectivity and reaction efficiency in complex biological tissues such as the brain. We believe that our findings will contribute to the progress in the newly emerging field of *in vivo* chemistry/chemical biology.

The vital roles of proteases in brain functions have not been fully characterized in molecular detail to date. Many issues remain to be elucidated, such as the existence/localization not only of proteases released inside the brain but also of proteases transported from outside the brain.^39^ The functional analysis of proteases using genetic techniques can be hindered in the complex situations in which there are a large number of genes for a target protease and when the expression and substrate patterns of related proteases are heavily overlapped. Our chemical KI method is expected to be complimentary to these genetic methods, and can enable the installation of a set of peptides as substrates for a target receptor, which will provide the ability to monitor diverse protease/peptidase activity in a live brain, potentially leading to the elucidation of the detailed functions of extracellular proteolysis in the brain. *In situ* zymography, a method for detecting protease activity in tissue sections, has been used to analyze the activity of MMPs.^40,41^ However, this method uses isolated brain slices instead of the whole live brain and has several limitations, including artifacts because of the postmortem brains used, the rather poor protease selectivity, and the low spatial resolution of the images.^40^ The method developed in the present study can be used to monitor protease activity in a living animal, which should provide results more indicative of the natural physiological state. The developed method also has high protease species selectivity because of the simplicity of synthesizing the desired peptide sequences by solid-phase-peptide synthesis, and the ability to monitor protease activity in proximity to a target receptor. In addition to the recording of the activity of a target protease in the brain as demonstrated herein, we envisage that our chemical KI strategy may be extended to the high-resolution mapping of important biological environmental changes (e.g., pH, metal ions, and bioactive small molecules) proximal to target receptors in live brains using various supramolecular and/or chemical biology-based sensors that have been developed to date.

## Methods

### Synthesis

All synthesis procedures and characterizations are described in the Supplementary Information.

### General methods for biochemical and biological experiments

SDS-PAGE and western blotting (WB) were carried out using a BIO-RAD Mini-Protean III electrophoresis apparatus. Samples were applied to SDS-PAGE and electrotransferred onto polyvinylidene fluoride membranes (BIO-RAD), followed by blocking with 5% nonfat dry milk in Tris-buffered saline containing 0.05% Tween 20. Primary antibody was indicated in each experimental procedure, and anti-rabbit IgG-HRP conjugate (CST, 7074S, 1:3,000) was utilized as the secondary antibody. Chemiluminescent signals generated with ECL Prime (Cytiva) were detected with a Fusion Solo S imaging system (Vilber Lourmat).

### General information for fluorescence imaging experiments

Fluorescence imaging was performed using a CLSM (Leica microsystems, Germany, TCS SP-8 or Carl Zeiss, Germany, LSM-800) equipped with 5× objective (numerical aperture (NA) = 0.25 dry objective for LSM-800), 10× objective (NA = 0.40 dry objective for TCP SP-8), 20× objective (NA = 0.75 oil objective for TCP SP-8), 40× objective (NA = 1.30 oil objective for TCP SP-8), 63× objective (NA = 1.40 oil objective for LSM-800), 100× objective (NA = 1.40 oil objective for TCP SP-8), and GaAsP detector. The excitation laser was derived from a 405 nm, 488 nm, 561 nm, 640 nm diode laser or a white laser and was set to an appropriate wavelength depending on the dye. Lightning deconvolution process (LAS X 3.5.5, Leica microsystems, Germany) was used in Supplementary Figure 7. Airyscan mode (Zeiss, Germany) was used in Figure 5D and 5G.

### Animal experiments

C57BL6/N mice were purchased from Japan SLC, Inc (Shizuoka, Japan). The animals were housed in a controlled environment (23 °C, 12 h light/dark cycle) and had free access to food and water, according to the regulations of the Guidance for Proper Conduct of Animal Experiments by the Ministry of Education, Culture, Sports, Science, and Technology of Japan. All experimental procedures were performed in accordance with the National Institute of Health Guide for the Care and Use of Laboratory Animals, and were approved by the Institutional Animal Use Committees of Kyoto University. Experimental details for injection of the labeling reagents are described in the Supplementary Information.

### Construction of AMPARs-based MMP9 sensor on HEK293T cells. (Figure 4)

HEK293T cells transfected with GluA2 were treated with 1 μM anchoring reagents **2(Ax555)** in the DMEM Glutamax including 10 mM HEPES at 17°C for 4 h. After removal of the medium, the cells were washed with HBSS (+) (Nacalai Tesque) twice and treated with 1 μM **MMP9-Leu** or **MMP9-DLeu** in the DMEM Glutamax including 10 mM HEPES at 17°C for 30 min. After removal of the reaction medium, the cells were washed with HBSS (+) twice and replaced with the DMEM Glutamax including 10 mM HEPES. Collagenase (Collagenase Type A, Fujifilm Wako, 034-24563, 1 mg/mL HBSS (+), 100 µL) was added to labeled cells on a microscope stage for CLSM observation. Confocal live imaging of labeled AMPA receptors were performed using a CLSM (TCS SP-8, Leica microsystems) equipped with a 40× objective and GaAsP detector. Fluorescence images were acquired using the 561 nm excitation for Ax555 and the 633 nm excitation for Ax647 derived from a white light laser.

### Construction of AMPARs-based MMP9 sensor in the live mouse brain. (Figure 5)

At 12 h after the LV injection of anchoring reagent **2(Ax555)** (100 µM, 4.2 µL), MeTz-peptide probe (100 µM, 4.2 µL) was injected into the LV. At 3, 6, and 9 h after the injection, the mouse was sacrificed under the deep isoflurane anesthesia or transcardially perfused with ice-cold 4% PFA/PBS(–) (pH 7.4). Brain sample was processed as described above and subjected to in gel/WB and CLSM analysis.

## Acknowledgments

The authors thank Ms. Tomoko Gonda, Ms. Yuka Nabeta, and Ms. Kumiko Nishizawa for technical support of biological experiments. The authors also thank Victoria Muir, PhD, from Edanz (https://jp.edanz.com/ac) for editing a draft of this manuscript. This work was supported by the Japan Science and Technology Agency (JST) ERATO (grant number JPMJER1802 to I.H.) and MEXT/JSPS KAKENHI Grant-in-Aid for Specially Promoted Research (grant number 23H05405 to I.H.), Scientific Research (B) (grant number 24K01627 to H.N.), Scientific Research (C) (grant number 23K04960 to S.S.), and JSPS Fellows (grant number 21J15773 to K.S.).

## Contributions

H.N. and I.H. initiated and designed the project. S.S., K.S., M.W., M.I., and H.N. performed synthesis and chemical labeling experiments in HEK293T cells and brains. H.N., S.S, K.S., and I.H. prepared the manuscript with contributions from the other authors.

## Corresponding author

Correspondence to Hiroshi Nonaka and Itaru Hamachi.

