## Supplementary Information for "In-brain construction of receptor-based protease sensors by coupling ligand-directed chemistry and click chemistry"

### Supplementary Figures

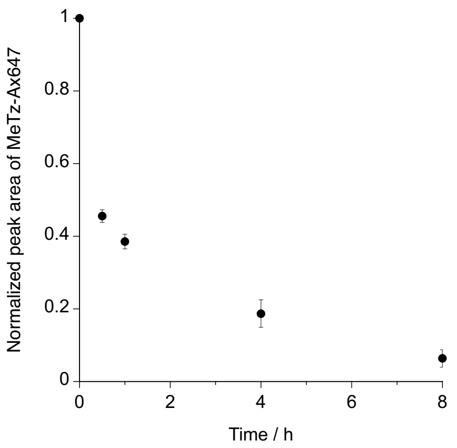


**Supplementary Figure 1 | Time course analysis of MeTz-Ax647 amount in the mouse brain after lateral ventricle (LV) injection of MeTz-Ax647.** MeTz-Ax647 (100 μM, 4.5 μL) in PBS (–) was injected into the mouse LV. After incubation to each time points, the whole brain was isolated and homogenated. The homogenate was extracted with 80% methanol/$H_2O$ (vol./vol.). After centrifugation, the supernatant was analyzed by RP-HPLC. RP-HPLC analysis was conducted on a Shimadzu Nexera system equipped with a fluorescent detector (630/650 nm (Ex/Em)) with a linear gradient of 0-40 % $CH_3CN$/10 mM triethylammonium acetate (TEAA) aq. (40 min). Data are presented as mean ± SEM.

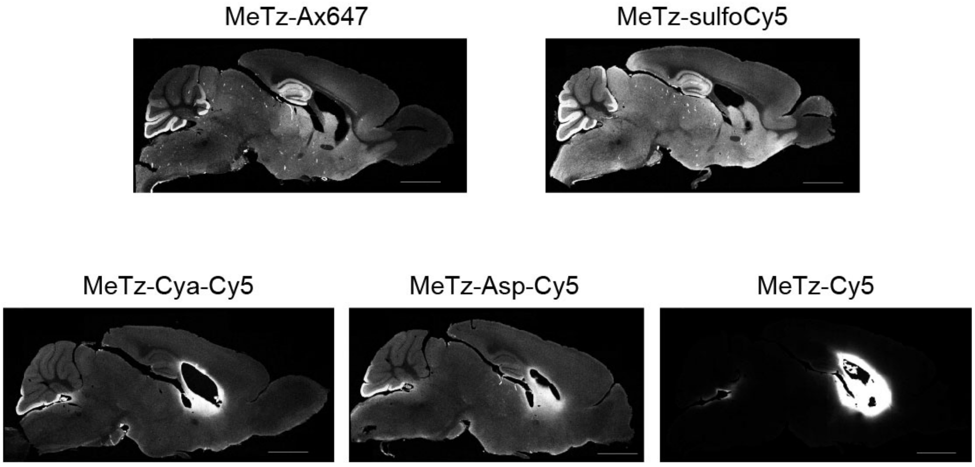


**Supplementary Figure 2 | CLSM imaging of saggital brain sections labeled with MeTz-Dyes.** At 24 h after the injection of **anchoring reagent 2(Ax555)** (100 μM, 4.5 μL), MeTz-dye (100 μM, 4.5 μL) was injected into the mouse LV. At 24 h after the injection, the mice were transcardially perfused with ice-cold 4% PFA/PBS(–) (pH 7.4). Brain slices were analyzed by a CLSM (TCS SP-8, Leica microsystems).

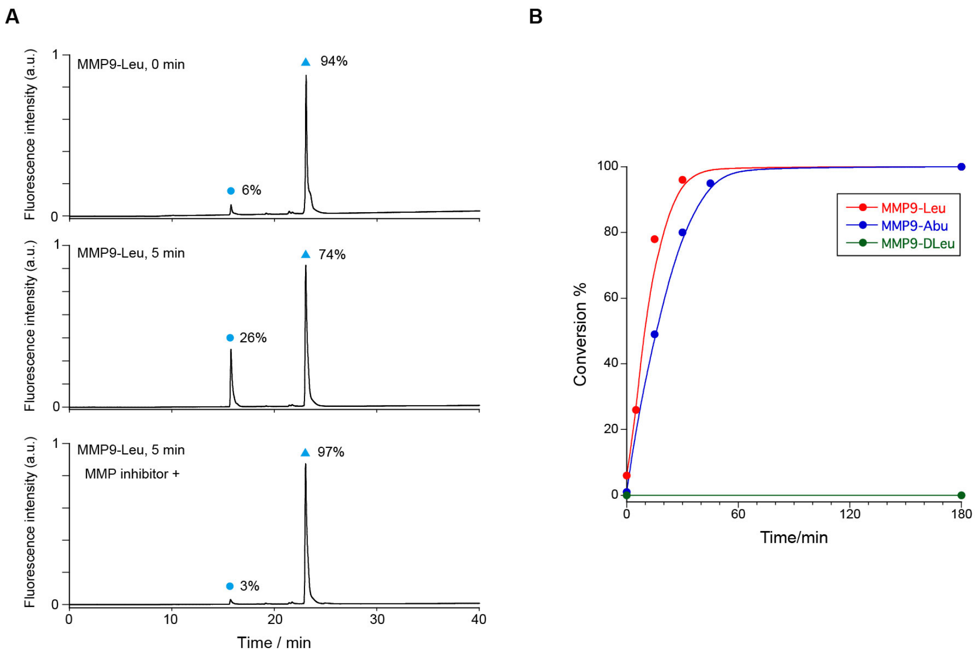


**Supplementary Figure 3 | HPLC analysis of MMP9-probes reactivity with a collagenase.**

A. Inhibition of MMP enzymatic reaction by pan-MMP inhibitor (ilomastat, 5 mM). Cyan circles indicate products and cyan triangles indicate starting materials. B. Time course analysis of reaction kinetics of each probe after the addition of a collagenase. A MMP9 probes (1 µM) were incubated in a collagenase (2.73 µg/mL in PBS (–), 50 µL) at 37 °C. After the addition of TEAA buffer (10 mM, 400 µL), reaction solution was filtered and analyzed by RP-HPLC (COSMOSIL 5C18-AR-II, 4.6 × 150 mm, mobile phase; $CH_3CN$ : 10 mM TEAA buffer (pH 7.0) = 0 : 100 → 50 : 50 (linear gradient over 40 min), flow rate = 1 mL/min, detection; fluorescence of Alexa Fluor 647, excitation = 633 nm and emission = 650 nm).

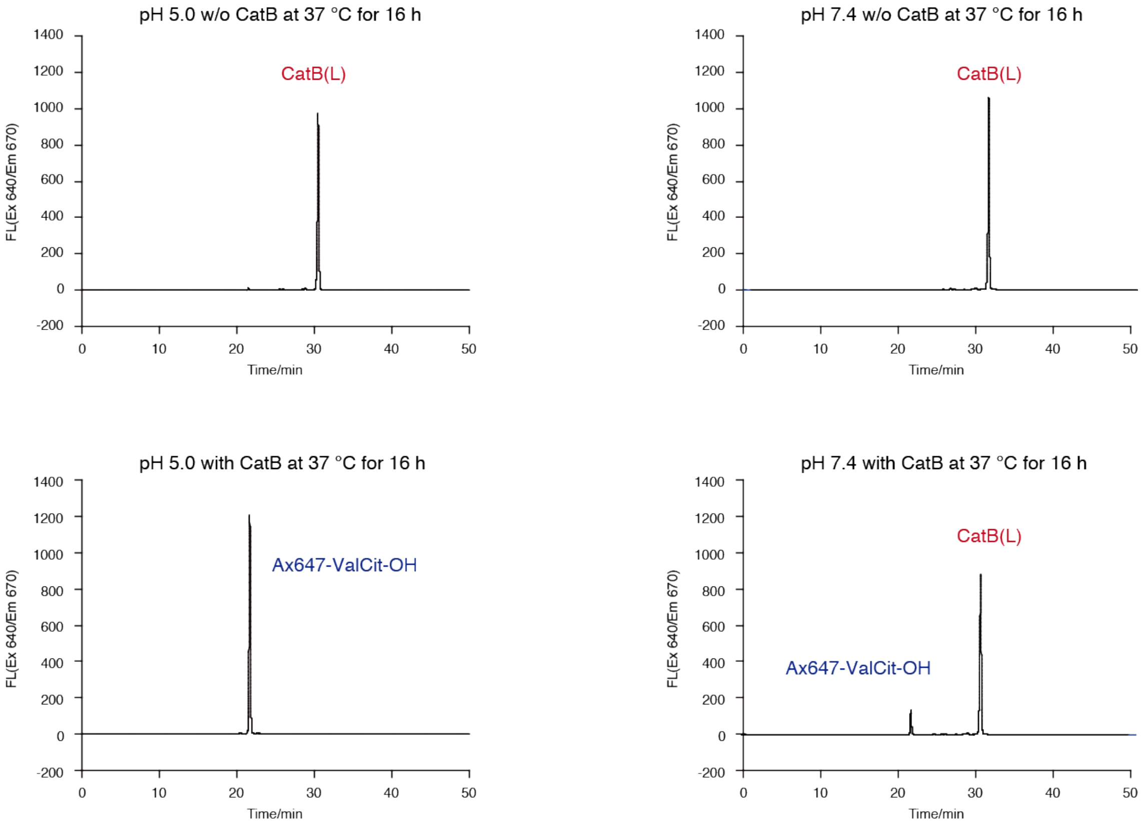


**Supplementary Figure 4 | Enzymatic reaction analysis of CatB probes in the buffer (pH 5.0 or pH 7.4).** A CatB(L) probe (10 μM) was incubated in a 20 mM NaOAc buffer (pH 5.0) or HBS (pH 7.4) containing 5 mM Cys (total volume = 200 μM) at 37 °C for 16 h in the presence or absence of cathepsin B (0.1 u, bovine spleen, SIGMA c6286). After heat inactivation at 95 °C for 3 min, 10 μL of reaction solution was analyzed by RP-HPLC (YMC Triart C18, 4.6 x 250 mm, mobile phase; $CH_3CN$ : 10 mM TEA AcOH buffer (pH 7.0) = 0 : 100 → 50 : 50 (linear gradient over 50 min), flow rate = 1 mL/min, temperature of column oven = 40 °C, detection; fluorescence of Alexa Fluor 647, excitation = 640 nm and emission = 670 nm).

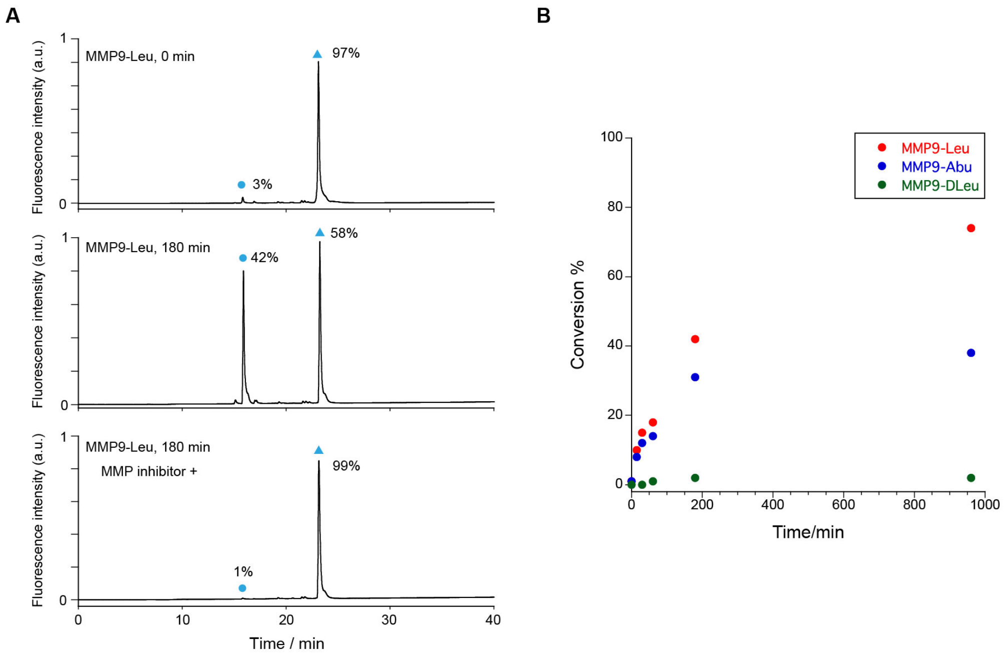


**Supplementary Figure 5 | HPLC analysis of MMP9-probe reactivity with the brain homogenate.** A. Inhibition of MMP enzymatic reaction by pan-MMP inhibitor (ilomastat, 1 mM). Red circles indicate products and red triangles indicate starting materials. B. Time course analysis of reaction kinetics of each probe after addition of the brain homogenate. A MMP9 probes (1 μM) were incubated in a hippocampus homogenate (1.15 mg/mL in PBS(–), 50 μL) at 37 °C. After the addition of TEAA buffer (10 mM, 400 μL), reaction solution was filtered and analyzed by RP-HPLC (COSMOSIL 5C18-AR-II, 4.6 × 150 mm, mobile phase; $CH_3CN$ : 10 mM TEAA buffer (pH 7.0) = 0 : 100 → 50 : 50 (linear gradient over 40 min), flow rate = 1 mL/min, detection; fluorescence of Alexa Fluor 647, excitation = 633 nm and emission = 650 nm).

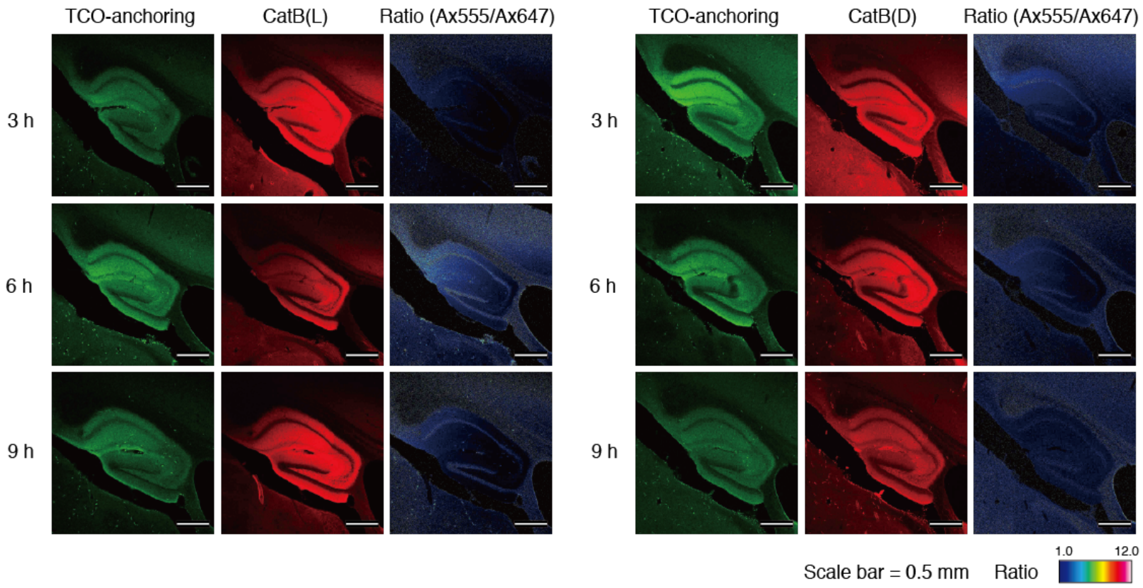


**Supplementary Figure 6 | Detection of CatB activity in the hippocampus area by CLSM.** 5× obejective, scale bar 0.5 mm. At 12 h after the LV injection of anchoring reagent **2(Ax555)** (100 μM, 4.2 μL), MeTz-peptide probe (100 μM, 4.2 μL) was injected into the LV. At 3, 6, 9 h after the injection, the mouse was sacrificed under the deep isoflurane anesthesia or transcardially perfused with ice-cold 4% PFA/PBS(–) (pH 7.4). Brain sample was processed as described above and subjected to in gel/WB and CLSM analysis.There was little change in the fluorescence images of samples labeled with the CatB(L) probe and those labeled with the CatB(D) probe, which suggested that the CatB activity was low in the extracellular region of the hippocampus.

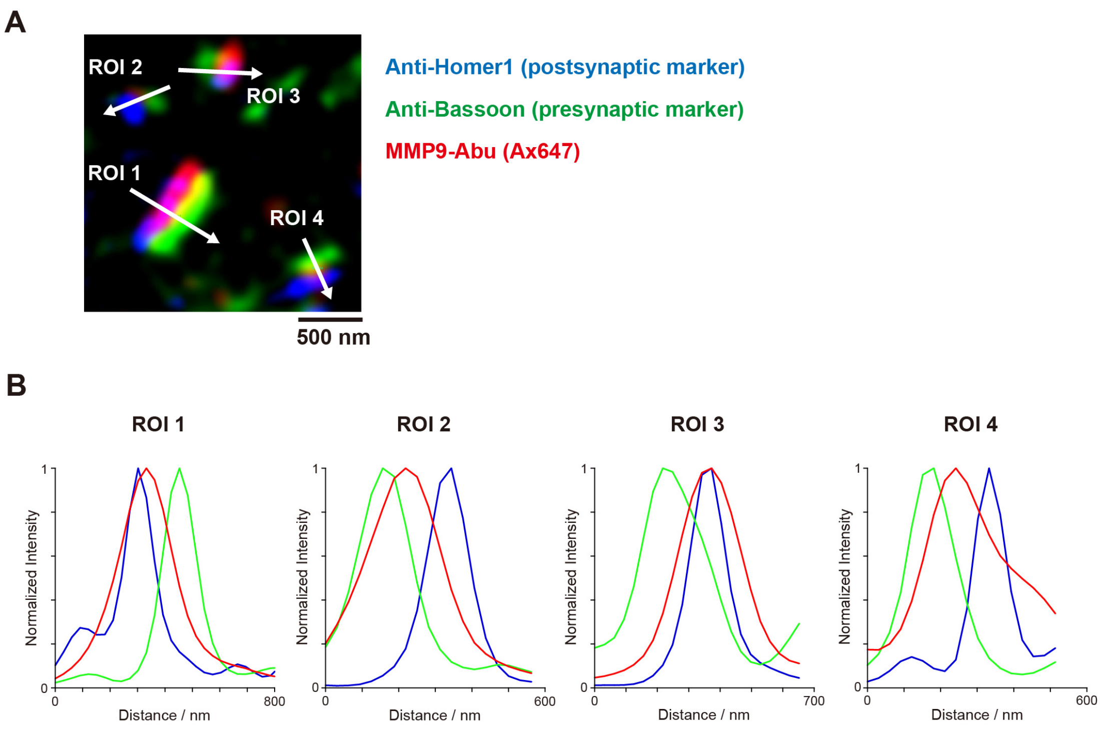


**Supplementary Figure 7 | High-resolution imaging of the brain section on which the MMP9-Abu tethered AMPAR sensor was constructed.** A. The cortical section was co-immunostained with presynaptic and postsynaptic markers, anti-Bassooon and anti-Homer1. B. Line plot analysis. Blue: Homer1, Red: Labeled AMPAR, Green: Bassoon. The data indicate that an AMPA receptor-based MMP9 sensor is indeed constructed within the synaptic cleft.

### Supplementary Methods

**Expression of receptors in HEK293T cells.**

HEK293T cells (ATCC) were cultured in Dulbecco's modified Eagle's medium (DMEM) supplemented with 10% fetal bovine serum (Sigma Aldrich), penicillin (100 units $mL^{-1}$), streptomycin (100 μg $mL^{-1}$) and amphotericin B (250 ng $mL^{-1}$) and incubated in a 5% $CO_2$ humidified chamber at 37 °C. For expression of each receptor, HEK293T cells ($2.0 \times 10^5$ cells) plated on a 3.5-cm dish (Corning) were transfected with a plasmid encoding rat GluA2 (GluA2flip(Q))[S1] or the control vector pCAGGS (kindly provided by Dr. H. Niwa from RIKEN) using Lipofectamine 2000 (Invitrogen) according to the manufacturer's instructions. The transfected cells were subjected to labeling experiments after 36–48 h of the transfection.

**Chemical labeling of AMPA receptors (AMPARs) in HEK293T cells. (Fig. 2)**

In the evaluation of anchoring efficiency for AMPARs, HEK293T cells transfected with GluA2 were treated with 1 μM **anchoring reagents 1-5** in the DMEM Glutamax at 37ºC for 3 h.

In the case of MeTz derivatives labeling, HEK293T cells transfected with GluA2 were treated with 1 μM **anchoring reagents 2(Ax555)** in the DMEM Glutamax at 37ºC for 3 h. The medium was removed and treated with 1 μM **MeTz derivatives** in the DMEM Glutamax at 37ºC for 1 h.

For WB analyses of labeled receptors, labeled cells were washed three times with PBS, lysed with radio immunoprecipitation assay (RIPA) buffer containing 1% protease inhibitor cocktail set III (Millipore, 539134), and mixed with 5× Laemmli sample buffer containing 250 mM DTT. In the experimental condition using anchoring reagent **5**, the lysate was reacted with 1 μM tetrazine-Ax647 (Tz-Ax647) for 5 min at room temperature. To quench excess Tz-Ax647, 10 μM TCO-PEG3-COOH was added to the lysate. The resulting lysate was mixed with 5× Laemmli sample buffer containing 250 mM DTT and subjected to SDS PAGE and WB analyses. SDS PAGE and WB analyses were performed as described in "General methods for biochemical and biological experiments". GluA2 was detected using a Anti-Ionotropic Glutamate receptor 2 antibody (abcam, ab206293, 1:3,000).

**Injection of reagents into the mouse LV. (Figures 3 and 5)**

Experiments were conducted according to the literature[S2,S3] using 5 weeks old mice (male, C57BL/6N strain; body weight 18–23 g). Under the deep anesthesia, the labeling reagent solution (4.5 μL) was directly injected into the LV using a microinjector (Nanoliter 2010, world precision instruments) (600 nL/min).

**Perfusion fixation and brain slices preparation. (Figures 3 and 5)**

Experiments were performed as described in the literature.[S4] Briefly, under deep isoflurane anesthesia, mice were transcardially perfused with ice-cold 4% PFA/PBS(–) (pH 7.4) or ice cold 9% glyoxal/8% acetic acid (pH 4.0). The isolated mouse brain samples were fixed overnight at 4 °C. After washing with PBS(–) (×3), the brain samples were immersed in 30% sucrose/PBS(–). Brain slices were prepared using a cryostat (Leica, CM-1950).

**Chemical labeling of endogenous AMPA receptors with LV injection. (Figures 3 and 5)**

For in gel/WB analysis, the mouse was sacrificed under the deep isoflurane anesthesia. The brain was isolated, washed twice with PBS, and lysed with RIPA buffer containing 1% protease inhibitor cocktail set III (Millipore, 539134). For brain lysates injected with anchoring reagent **5**, the lysates were reacted with 1 μM Tz-Ax647 for 5 min at room temperature. After that, 10 μM TCO-PEG3-COOH was added to quench excess Tz-Ax647. The obtained lysates were mixed with 5× Laemmli sample buffer containing 250 mM DTT.

In the case of kinetic analysis, the mice were transcardially perfused with PBS(–) containing 20 μM of TCO-PEG$_3$-COOH. then brain isolated and lysed with RIPA buffer containing 1% protease inhibitor cocktail set III (Millipore, 539134) and 100 μM of TCO-PEG$_3$-COOH.

The lysate was incubated at 4 °C for 30 min and centrifuged at 4 °C and 12,000 rpm for 10 min. After mixing with 5× Laemmli sample buffer containing 250 mM DTT, SDS-PAGE and WB analyses were performed as described in "General methods for biochemical and biological experiments". The Ax647-labeled AMPAR was detected by Ax647-fluorescence. The GluA2 was detected using a rabbit anti-GluA2 antibody (abcam, ab206293, 1:3,000).

For CLSM analyses, the mouse was transcardially perfused with ice cold 4% PFA/PBS(–) or ice cold 9% glyoxal/8% acetic acid (pH 4.0)[S5] at 24 h after the injection. The brain was isolated and sectioned by cryostat. Imaging was performed using a CLSM.

**Immunohistochemical staining for GluA2 (Figures 2E and 2F) and Homer1/Bassoon (Supplementary Figure 7).**

The brain slices (50 μm thickness) were permeabilized with PBS(–) containing 0.1% triton X-100 for 15 min and blocked with 10% normal goat serum (NGS) in PBS(–) containing 0.1% triton X-100 for 30 min. Then, primary antibody reaction was conducted with the following antibodies in PBS(–) containing 0.1% triton X-100 at 4 °C overnight. Secondary antibody reaction was conducted with appropriate antibodies in PBS(–) containing 0.1% triton X-100 at r.t. for 1 h.

For GluA2 staining, the mouse was transcardially perfused with ice cold 4% PFA/PBS(–) or ice cold 9% glyoxal/8% acetic acid (pH 4.0). A rabbit anti-GluR2 (Genetex, GTX66722, 1:500) was used as a primary antibody. Secondary antibody reaction was conducted with a goat anti-rabbit IgG H&L (Alexa Fluor® 555) (abcam, ab150078, 1:500).

For Homer1 and Bassoon staining, the mouse was transcardially perfused with ice cold 4% PFA/PBS(–). A mouse anti-Homer1 (abcam, ab184955, 1:1000) and a rabbit anti-Bassoon (abcam, ab82958, 1:1000) were used as primary antibodies. Secondary antibody reaction was conducted with a goat anti-mouse IgG H&L (Alexa Fluor® 488) (abcam, ab150113, 1:200) and a goat anti-rabbit IgG H&L (Alexa Fluor® 405) (abcam, ab175652, 1:200).

**Tissue clearing of mouse brain with CUBIC protocol. (Figure 3G)**

Experiments were conducted according to the literature.[S6] In brief, the **MeTz-Ax647** labeled and PFA-fixed brain was delipidated with CUBIC-L, cleared with CUBIC-R, and embedded in CUBIC-R solution. The 2D macro-imaging was performed with a gel imager (VILBER, France, Fusion). The 3D imaging was performed with CLSM (Leica microsystems, Germany, TCS SP-8).

#### Synthesis and Characterization of Compounds

##### General materials and methods for organic synthesis

All chemical reagents and solvents were obtained from commercial suppliers (SIGMA-Aldrich, Tokyo Chemical Industry (TCI), FUJIFILM Wako Pure Chemical Corporation, or Watanabe Chemical Industries) and used without further purification. Thin layer chromatography (TLC) was performed on silica gel 60 $F_{254}$ precoated aluminum sheets and glass plate (Merck) and visualized by fluorescence quenching or ninhydrin staining. Chromatographic purification was conducted by flash column chromatography on silica gel 60N (neutral, 40–50 µm, Kanto Chemical) or Biotage isolera system equipped with a Biotage® Sfär cartridge. $^1$H NMR spectra were recorded in deuterated solvents on a JEOL ECS400 (400 MHz) or JEOL ECZ600R (600 MHz). Chemical shifts were referenced to residual solvent peaks or tetramethylsilane ($\delta = 0$ ppm). Multiplicities are abbreviated as follows: *s* = singlet, *d* = doublet, *t* = triplet, *q* = quartet, *quin* = quintet, *m* = multiplet. MALDI-TOF Mass spectra were measured on UltrafleXtreme (Bruker Daltonics). High resolution mass spectra were measured on an Exactive (Thermo Scientific) equipped with electron spray ionization (ESI). Reversed-phase HPLC (RP-HPLC) was carried out on a Hitachi Chromaster system equipped with a diode array using a Cosmosil 5C18AR2 (Nacalai tesque) and a YMC-Pack ODS-A column (YMC Co. Ltd.).

Abbreviation

Ax647: AlexaFluor® 647

Ax555: AlexaFluor® 555

Boc: tert-butoxycarbonyl

DCM: Dichloromethane

DIPEA: *N*,*N*-Diisopropylethylamine

DMF: *N*,*N*-dimethylformamide

DMSO: dimethyl sulfoxide

DSC: *N*,*N'*-disuccinimidyl carbonate

EDCI: 1-ethyl-3-(3-dimethylaminopropyl)carbodiimide

EtOAc: Ethyl acetate

HBTU: 1-[Bis(dimethylamino)methylene]-1H-benzotriazolium 3-Oxide Hexafluorophosphate

HOBt: 1-hydroxybenzotriazole

NHS: *N*-hydroxysuccinimide

TCO: *trans*-cyclooctene

TEA: Triethylamine

TFA: Trifluoroacetic acid

TSTU: *N*,*N*,*N'*,*N'*-Tetramethyl-*O*-(N-succinimidyl)uronium Tetrafluoroborate

TEAA: triethylammonium acetate

MeTz-$NH_2$: Methyltetrazine-amine

**Synthesis of 2(Ax647)**

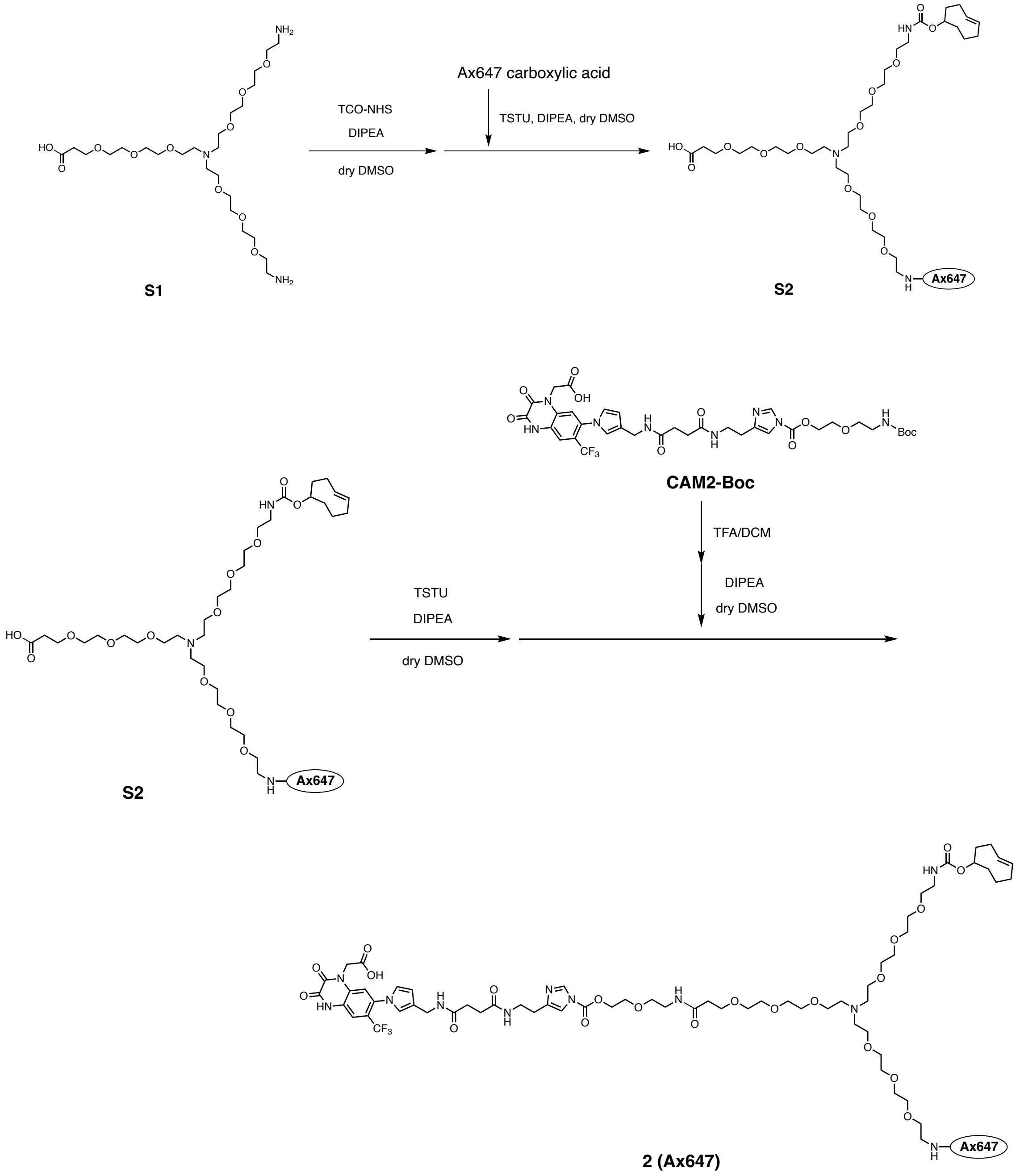


Compound **S2**: To a solution of **S1** (8.4 mg, 14.7 μmol) in dry DMSO (500 μL), TCO-NHS ester (6.3, 23.5 μmol) in dry DMSO (420 μL) and DIPEA (27.6 μL, 158 μmol) were added. The mixture was stirred at room temperature for 2 h under Ar atmosphere (Solution A). The mixture of Alexa Fluor™ 647 carboxylic acid tris(triethylammonium) salts (10 mg, 7.91 μmol) in dry DMSO (500 μL), 20 μg/μL TSTU/DMSO solution (155 μL, 3.1 mg, 10.3 μmol), and DIPEA (6.9 μL 39.5 μmol) was stirred at room temperature for 3 h under Ar atmosphere (Solution B). These two solutions were mixed and the reaction mixture was stirred for 3 h at room temperature under Ar atmosphere. The reaction mixture was purified by RP-HPLC (column; Cosmosil 5C18AR2, 250 x 20 mm, mobile phase; $CH_3CN$ : 10 mM TEAA aq. = 10 : 90 → 32 : 68 (linear gradient over 30 min), flow rate; 9 mL/min, detection; UV absorbance at 220 nm) to give **7** as a blue solid (3.9 mg, 2.5 μmol, 32% yield). HR ESI-MS (calc. for $C_{70}H_{107}N_5O_{26}S_4$): $[M-2H]^{2-}$ = 780.8042 (calc. = 780.8049).

**2(Ax647)**: The compound **CAM2-Boc** was synthesized according to the previously reported method.[S1] To a solution of **CAM2-Boc** (2.70 mg, 3.35 μmol) in dry DCM (400 μL), TFA (100 μL mL) was added and the reaction mixture was stirred at room temperature for 1.5 h. After azeotropically removal of the solvent with $CH_3CN$:toluene = 1:1 (400 μL, ×3), the Boc-deprotected compound was dissolved in dry DMSO (500 μL) (Solution A). The mixture of compound **7** (1.3 mg, 0.83 μmol) in dry DMSO (400 μL), 10 μg/μL TSTU/DMSO solution (37.5 μL, 0.375 mg, 1.25 μmol), and DIPEA (5.0 μL 29 μmol) was stirred at room temperature for 2.5 h under Ar atmosphere (Solution B). These two solutions were mixed and the reaction mixture was stirred for 3 h at room temperature under Ar atmosphere. The reaction mixture was purified by RP-HPLC (column; Cosmosil 5C18AR2, 250 x 20 mm, mobile phase; $CH_3CN$ : 10 mM TEAA aq. = 5 : 95 → 40 : 60 (linear gradient over 40 min), flow rate; 9 mL/min, detection; UV absorbance at 220 nm) to give **2(Ax647)** as a blue solid (0.13 μmol, 16% yield, determined by measurement of UV-absorbance). HR ESI-MS (calc. for $C_{100}H_{137}F_3N_{13}O_{34}S_4Na$): $[M-3H+Na]^{2-}$ = 1136.4085 (calc. = 1136.4084).

**Synthesis of CAM2-N-TCO-Ax555 (2(Ax555))**

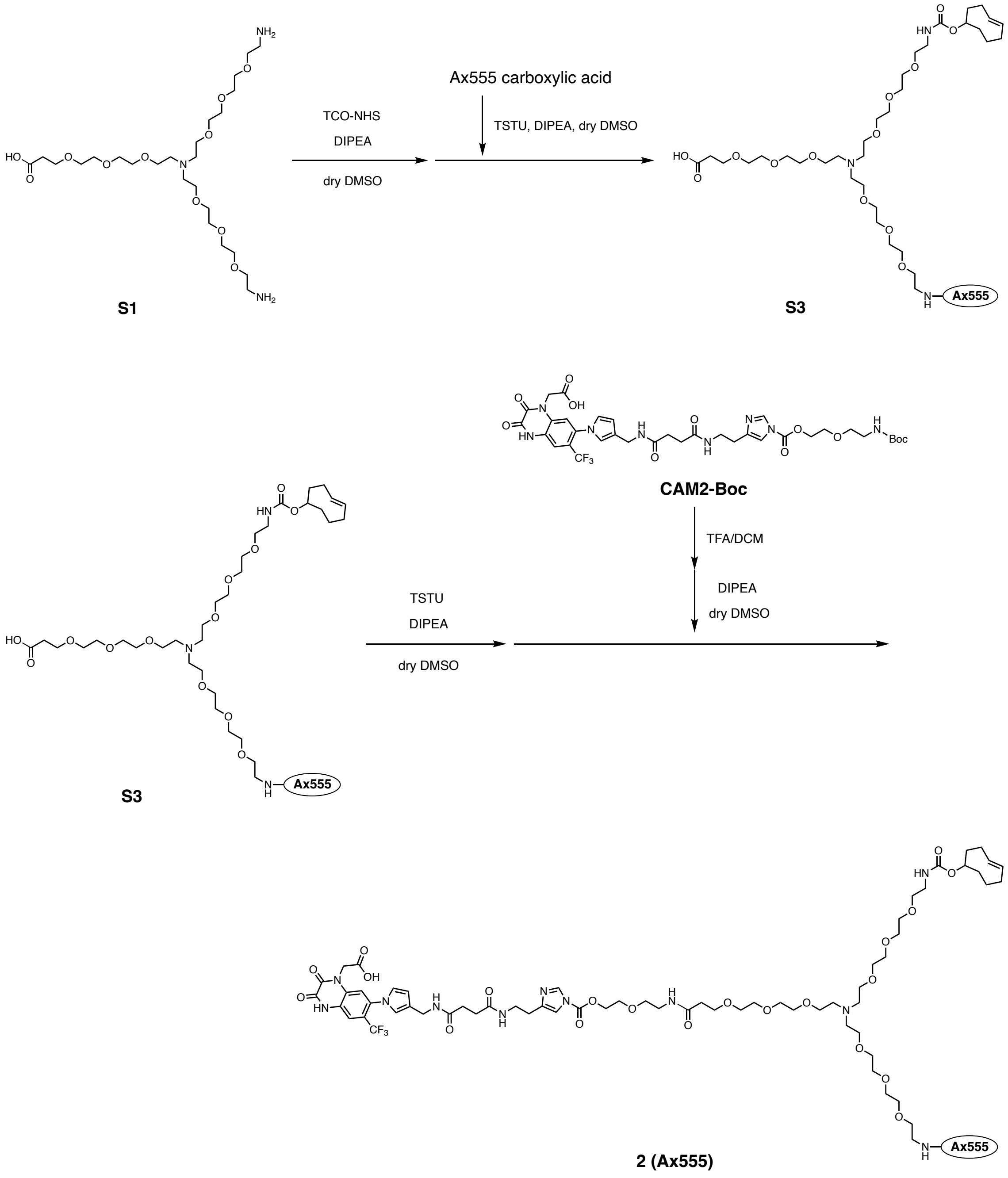


Compound **S3**: This compound was synthesized according to the similar way for the synthesis of S2. HR ESI-MS (calc. for $C_{68}H_{105}N_5O_{26}S_4$): $[M-2H]^{2-}$ = 767.7965 (calc. = 767.7971).

**2 (Ax555):** This compound was prepared from **CAM2-Boc** according to the similar way for the synthesis of **2 (Ax647)**. HR ESI-MS **2** (calc. for $C_{98}H_{136}F_3N_{13}O_{34}S_4$): $[M-2H]^{2-}$ = 1111.9069 (calc. = 1111.9079).

#### Synthesis of compound 3

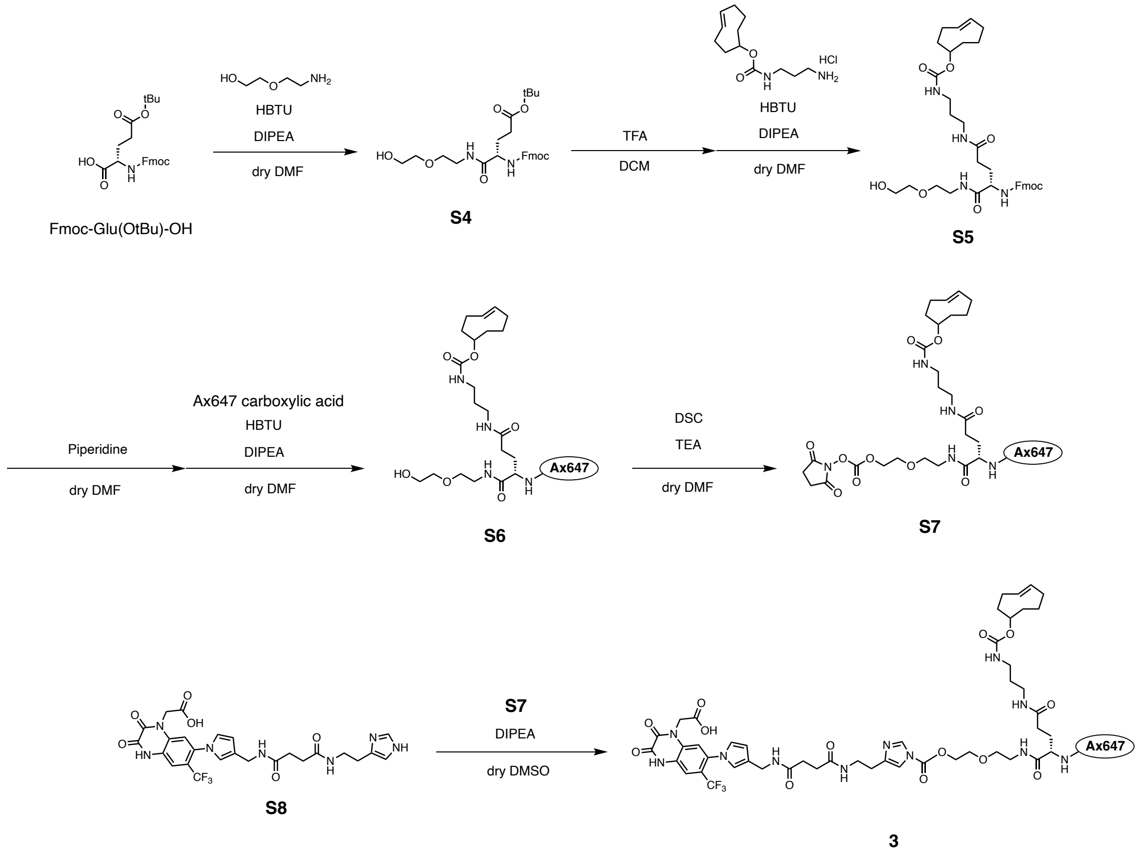


Compound **S4**: To a solution of Fmoc-Glu(O$^t$Bu)-OH (230 mg, 0.51 mmol) in dry DMF (2mL) was added 2-(2-aminoethoxy)ethanol (93 µL, 0.93 mmol), HBTU (235 mg, 0.62 mmol), DIPEA (222 µL, 1.28 mmol). The reaction mixture was stirred for 2.5 h at room temperature under $N_2$ atmosphere. Sat. $NaHCO_3$ was added and extracted with hexane:ethyl acetate =4:1 (50 mL × 3). The organic layer was dried over $Na_2SO_4$. After removal of the solvent by evaporation, the residue was purified by flash column chromatography on silica gel ($CHCl_3$ / MeOH = 100 : 0 to 90:10) to give **S4** (188 mg, 0.35 mmol, 70 % yield) as a colorless oil. $^1$H-NMR (400 MHz, $CDCl_3$) δ 7.79 (d , *J* = 7.2 Hz, 2H), 7.63 (d, *J* = 7.2 Hz, 2H), 7.43 (t, *J* = 7.6 Hz, 2H), 7.34 (t, *J* = 7.6 Hz, 2H), 6.06 (d, *J* = 8.0 Hz, 2H), 4.42-4.40 (m, *J* = 7.6 Hz, 2H), 4.24 (q, *J* =6.8 Hz, 2H), 3.73-3.72 (m, 2H), 3.59-3.58 (m, 4H), 3.50 (t, *J* = 4.8 Hz, 2H), 2.47-2.33 (m, 2H), 2.15-2.10 (m, 1H), 2.01-1.96 (m, 1H), 1.49 (s, 9H).

Compound **S5**: To a solution of **S4** (14 mg, 27 µmol) in DCM (0.9 mL) was added TFA (100 µL). The reaction mixture was stirred for 1 h at room temperature. After azeotropically removal of the solvent with toluene (1 mL × 3), the residue was dissolved in dry DMF (1 mL). TCO-amine (7.0 mg, 27 μmol), HBTU (15 mg, 40 μmol), and DIPEA (14 μL, 81 μmol) were added and the reaction mixture was stirred for 4 h at room temperature. After removal of the solvent by evaporation, the residue was purified by flash column chromatography on silica gel ($CHCl_3$ / MeOH = 100 : 0 to 90:10) to give **S5** (13 mg, 20 μmol, 72 % yield, including impurities) as a colorless oil. $^1$H-NMR (400 MHz, $CDCl_3$) δ 7.76 (d , *J* = 7.6 Hz, 2H), 7.59 (d, *J* = 6.0 Hz, 2H), 7.39 (t, *J* = 7.6 Hz, 2H), 7.30 (t, *J* = 7.6 Hz, 2H), 6.20 (d, *J* = 7.2 Hz, 2H), 5.64-5.42 (m, 2H), 4.33 (d, *J* = 7.2 Hz, 2H), 4.30-4.26 (m, 2H), 4.19 (t, *J* = 6.8 Hz, 2H), 3.73-3.71 (m, 2H), 3.59-3.58 (m, 4H), 3.50-3.48 (m, 2H), 3.37-3.32 (m, 1H), 3.24-3.15 (m, 4H), 2.35 -2.29 (m, 2H), 1.94-1.87 (m, 2H), 1.74-1.64 (m, 2H), 1.51-1.40 (m, 10H).

Compound **S7**: **S5** (8.7 mg, 13 μmol) was dissolved in 5% piperazine/DMF solution (500 μL). The reaction mixture was stirred for 1 h at room temperature. After azeotropically removal of the solvent with toluene (1 mL × 3), the residue was dissolved in dry DMF (500 μL) and Alexa Fluor™ 647 carboxylic acid tris(triethylammonium) salts (10.3 mg, 11.9 μmol), HBTU (6.5 mg, 17 μmol), and DIPEA (6.7 mg, 39 μmol) were added. The reaction mixture was stirred for 12 h at room temperature. The residue was purified by RP-HPLC (column; Cosmosil 5C18AR2, 250 x 10 mm, mobile phase; $CH_3CN$ : 10 mM $AcONH_4$ aq. = 10 : 90 → 40 : 60 (linear gradient over 30 min), flow rate; 9 mL/min, detection; UV (220 nm)) to give **S6** (5.0 mg, 3.9 μmol) as a blue solid. MALDI-TOF-Mass *m/e* calcd for $[M-H]^-$ 1281.44, found 1281.98. To a solution of compound **S6** (5.0 mg, 3.89 μmol) dissolved in dry DMF (0.4 mL), DSC (9.9 mg, 38.9 μmol) and TEA (5.3 μmol, 38.9 μmol) were added. Aftter stirring at room temperature under Ar atmosphere for 6 h, the reaction mixture stored O/N at -80℃. Then DSC (9.9 mg, 38.9 μmol) and TEA (5.3 μmol, 38.9 μmol) were further added. Aftter stirring at room temperature under Ar atmosphere for 2 h, the reaction mixture was purified by RP-HPLC (Cosmosil 5C18AR2, 250 x 10 mm, mobile phase; $CH_3CN$ : 10 mM TEAA aq. = 10 : 90 → 45 : 55 (linear gradient over 35 min), flow rate; 9 mL/min, detection; UV absorbance at 650 nm) to give a compound **S7** as a blue solid (1.83 μmol, 47%). MALDI-TOF-Mass *m/e* calcd for $[M-H]^-$ 1422.45, found 1423.04. The product was used for the next step without further purification.

Compound **3**: To a solution of **S7** (2.78 μmol) in dry DMSO (400 μL) was added **S8** (1.87 μmol) in dry DMSO (20 μL) and DIPEA (5 μL, 28 μmol). The reaction mixture was stirred for O/N at room temperature under $N_2$ atmosphere. The reaction mixture was purified by RP-HPLC (column; Cosmosil 5C18AR2, 250 x 10 mm, mobile phase; $CH_3CN$ : 10 mM TEAA aq. = 10 : 90 → 45 : 55 (linear gradient over 35 min), flow rate; 9 mL/min, detection; UV (220 nm)) to give **3** (0.51 μmol, 27% yields) as a blue solid. HR ESI-MS (calc. for $C_{83}H_{101}F_3N_{13}O_{26}S_4Na$): $[M-3H+Na]^{2-}$ = 951.7870 (calc. = 951.7862).

**Synthesis of CAM2-Lys(TCO)-Ax647 (4)**

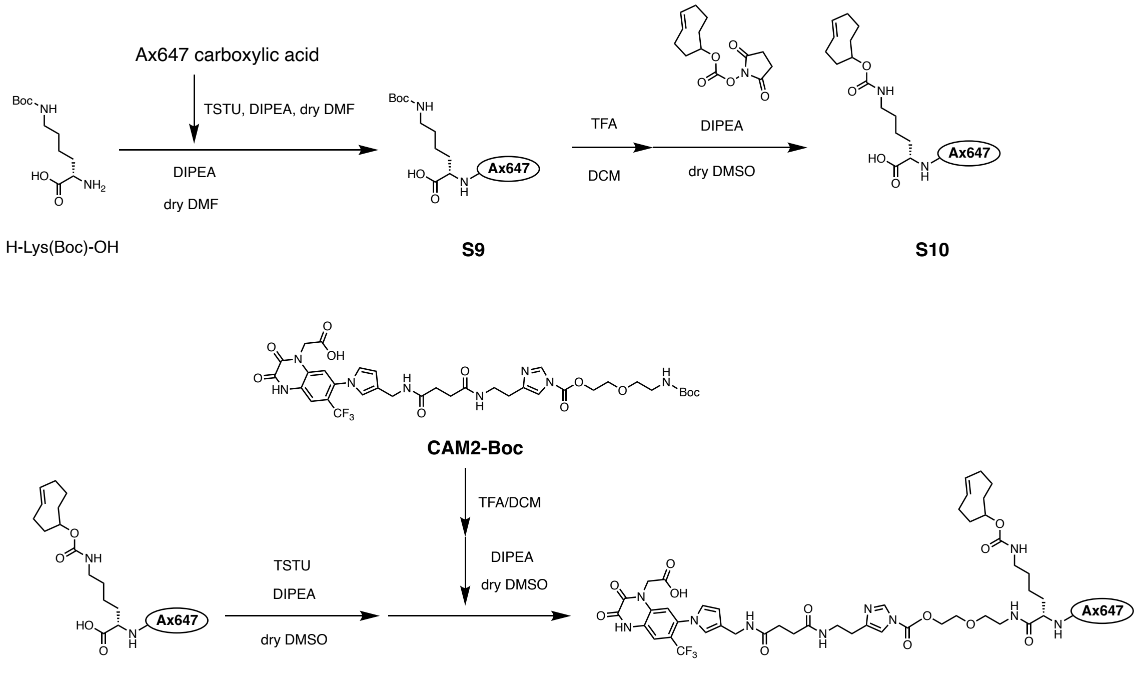


H-Lys(Boc)-OH S9 S10

CAM2-Boc

S10 4

Compound **S9**:

To a solution of Alexa Fluor™ 647 carboxylic acid tris(triethylammonium) salts (5 mg, 4.3 μmol) in dry DMF (300 μL), TSTU (1.62 mg, 5.4 μmol) in dry DMF (16.2 μL) and DIPEA (7.5 μL, 43 μmol) were added. The mixture was stirred at room temperature for 2 h under Ar atmosphere. H-Lys(Boc)-OH (6.2 mg, 25 μmol) and DIPEA (9.0 μL, 52 μmol) in dry DMF (200 μL) was added to the reaction mixture. After stirring for 12 h at room temperature under Ar atmosphere, the reaction mixture was purified by RP-HPLC (column; Cosmosil 5C18AR2, 250 x 10 mm, mobile phase; $CH_3CN$ : 10 mM ammonium acetate = 0 : 100 → 30 : 70 (linear gradient over 50 min), flow rate; 3 mL/min, detection; UV absorbance at 220 nm). After lyophilization of the target fraction, the product was dissolved with 100 mM TEAA (pH 7.0) and lyophilized to give **S9** as a blue solid. (4.2 mg, 3.1 µmol, 70% yield). HR ESI-MS (calc. for $C_{47}H_{63}N_4O_{17}S_4$): $[M-3H]^{3-}$ = 361.1031 (calc. = 361.1029).

Compound **4**:

To a solution of compound **S9** (5.60 mg, 6.07 µmol) in DCM (500 µL) was added TFA (100 µL) and the reaction mixture was stirred at room temperature for 2.5 h. After azeotropically removal of the solvent with $CH_3CN$:toluene = 1:1 (200 µL, ×3), the Boc-deprotected compound was dissolved in dry DMSO (500 µL). TCO-NHS (6.50 mg, 24.3 µmol) and DIPEA (33 µL, 189 µmol) was added to the solution. After stirring overnight under Ar atmosphere, the reaction mixture was purified by RP-HPLC (column; Cosmosil 5C18AR2, 250 x 20 mm, mobile phase; $CH_3CN$ : 10 mM TEAA aq. = 0 : 100 → 30 : 70 (linear gradient over 50 min), flow rate; 9 mL/min, detection; UV absorbance at 220 nm) to give **S10** as a blue solid (3.4 mg, 2.39 µmol as 3TEA salts, 39% yield). MALDI-MS (calc. for $C_{51}H_{69}N_4O_{17}S_4$): $[M-H]^{-}$ = 1137.8 (calc. = 1137.4). The product was used directly for the next step without further purification. The mixture of compound **S10** (3.4 mg, 2.39 µmol) in dry DMSO (500 µL), 21.2 µg/µL TSTU/DMSO solution (34 µL, 0.72 mg, 2.39 µmol), DIPEA (4.2 µL, 24.1 µmol) was stirred at room temperature under Ar atmosphere. After strring for 2 h, 21.2 µg/µL TSTU/DMSO solution (34 µL, 0.72 mg, 2.39 µmol) and DIPEA (4.2 µL, 24.1 µmol) were added to the reaction mixture and the mixture was stirred for 2 h (Solution A). To a solution of **CAM2-Boc** (5.80 mg, 7.19 µmol) in dry DCM (500 µL), TFA (100 µL mL) was added and the reaction mixture was stirred at room temperature for 1.5 h. After azeotropically removal of the solvent with $CH_3CN$:toluene = 1:1 (200 µL, ×3), the Boc-deprotected compound was dissolved in dry DMSO (600 µL) (Solution B). 200 µL of Solution B (2.40 µmol) was added to Solution A. After stirring for 1.5 h, 200 µL of Solution B (2.40 µmol) and DIPEA (4.2 µL, 24.1 µmol) were added. After stirring for 1.5 h, 200 µL of Solution B (2.40 µmol) and DIPEA (4.2 µL, 24.1 µmol) were added and the mixture was stirred overnight. The reaction mixture was purified by RP-HPLC (column; Cosmosil 5C18AR2, 250 x 10 mm, mobile phase; $CH_3CN$ : 10 mM TEAA aq. = 0 : 100 → 30 : 70 (linear gradient over 55 min), flow rate; 3 mL/min, detection; UV absorbance at 220 nm) to give **4** as a blue solid (0.32 µmol, 5% yield in 2 steps, determined by measurement of UV-absorbance). HR ESI-MS (calc. for $C_{81}H_{98}F_3N_{12}O_{25}S_4$): $[M-3H]^{3-}$ = 607.8543 (calc. = 607.8539).

**Synthesis of MeTz-Ax647**

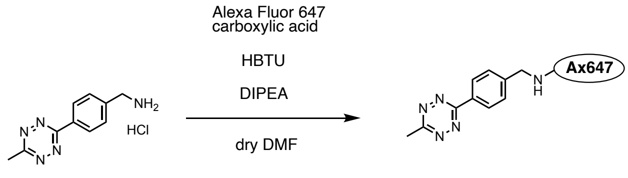


**MeTz-Ax647**

To a solution of Alexa Fluor™ 647 carboxylic acid tris(triethylammonium) salts (1.5 mg, 1.3 µmol) in dry DMF (500 µL), HBTU (4.0 mg, 10.5 µmol), MeTz-$NH_2$ hydrochloride (4.1 mg, 17 µmol), and DIPEA (4.1 µL, 23.5 µmol) were added. The mixture was stirred at room temperature overnight under Ar atmosphere. The reaction mixture was purified by RP-HPLC (column; Cosmosil 5C18AR2, 250 x 10 mm, mobile phase; $CH_3CN$ : 10 mM ammonium acetate aq. = 0 : 100 → 50 : 50 (linear gradient over 50 min), flow rate; 3 mL/min, detection; UV absorbance at 220 nm) to give **MeTz-Ax647** as a blue solid (0.792 µmol, 46% yield). HR ESI-MS (calc. for $C_{46}H_{52}N_7O_{13}S_4$): $[M\text{-}3H]^{3-}$ = 346.0848 (calc. = 346.0841).

**Synthesis of MeTz-Cya-Cy5**

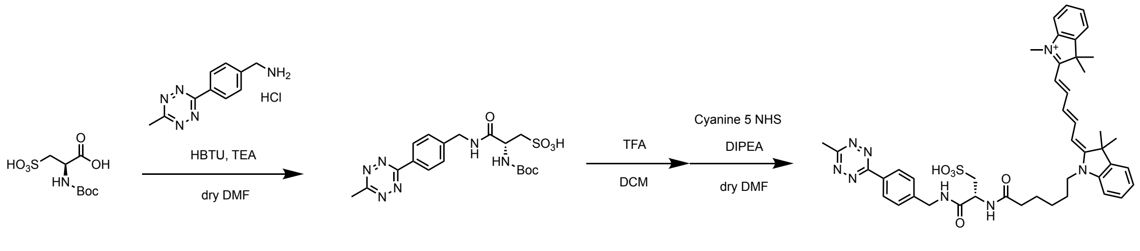


**S11** **MeTz-Cya-Cy5**

To a solution of Boc-Cya-OH (10.4 mg) in dry DMF (1mL) was added MeTz-$NH_2$・HCl (3.1 mg, 13 µmmol), HBTU (10.6 mg, 28 µmol), and $Et_3N$ (20.0 µL, 146 µmol). The reaction mixture was stirred for O/N at room temperature under $N_2$ atmosphere. After removal of the solvent by evaporation, the residue was purified by flash column chromatography on silica gel ($CHCl_3$ / MeOH = 100 : 0 to 90:10) to give **S11** (10.2 mg, including impurities) as a pink solid. $^1$H-NMR (400 MHz, $CD_3OD$) δ 8.43 (d , $J$ = 8.0 Hz, 2H), 7.54 (d, $J$ = 8.4 Hz, 2H), 4.59-4.45 (m, 3H), 3.29-3.28 (m, 2H), 3.00 (s, 3H) 1.43 (s, 9H). HR ESI-MS (calc. for $C_{18}H_{23}N_6O_6S$): $[M–H]^-$ = 451.1403 (calc. = 451.1405).The crude product was directly used for the next step without further purification. To solution of **S11** (2 mg) in $CH_2Cl_2$ (0.3 mL) was added TFA (150 µL). The reaction mixture was stirred for 3 h at room temperature. After azeotropically removal of the solvent with toluene (1 mL × 3), the residue was dissolved in dry DMF (200 µL) and **Cy5-NHS** in dry DMF

(100 μL), $Et_3N$ (10 μL) were added. The reaction mixture was stirred for O/N at room temperature and purified by RP-HPLC (column; Cosmosil 5C18AR2, 250 x 10 mm, mobile phase; $CH_3CN$ : 10 mM TEAA aq. = 50 : 50 → 80 : 20 (linear gradient over 30 min), flow rate; 9 mL/min, detection; UV (220 nm)) to give **MeTz-Cya-Cy5** (0.22 mg, 0.3 μmol) as a blue solid. HR ESI-MS (calc. for $C_{46}H_{53}N_8O_4$): $[M+H]^+$ = 781.4180 (calc. = 781.4184).

#### Synthesis of MeTz-Asp-Cy5

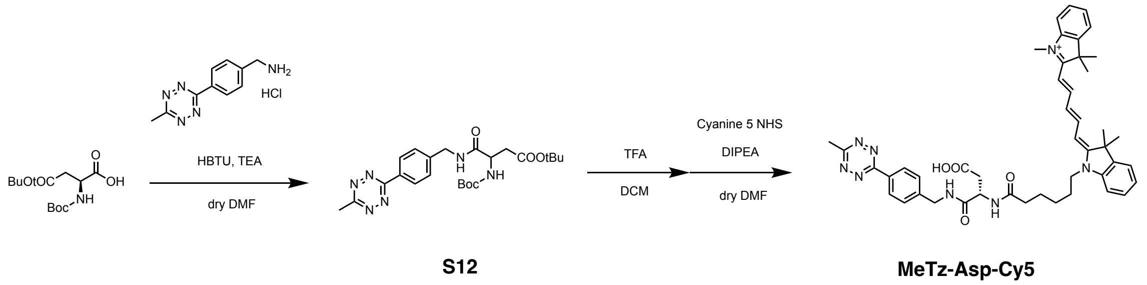


To a solution of Boc-Asp(OtBu)-OH (8.2 mg, 28 μmol) in dry DMF (1mL) was added MeTz-$NH_2$・HCl (3.2 mg, 13 μmmol), HBTU (10.8 mg, 28 μmol), $Et_3N$ (20 μL, 146 μmol). The reaction mixture was stirred for O/N at room temperature under $N_2$ atmosphere. After removal of the solvent by evaporation, the residue was dissoleved with $CHCl_3$. The solution was washed sat. $NaHCO_3$ (30 mL x2) and the organic layer was dried over $Na_2SO_4$. After removal of the solvent by evaporation, the residue was purified by flash column chromatography on silica gel ($CHCl_3$ / MeOH = 100 : 0 to 90:10) to give **S12** (9.2 mg; including impurities) as a pink solid. $^1$H-NMR (400 MHz, $CDCl_3$) δ 8.53(d , $J$ = 8.4 Hz, 2H), 7.47 (d, $J$ = 8.0 Hz, 2H), 4.64-4.51 (m, 3H), 3.09 (s, 3H), 2.99-2.62 (m, 2H),1.45-1.42 (m, 18H). The crude product was directly used for the next step without further purification. To solution of **S12** (2 mg) in $CH_2Cl_2$ (0.3 mL) was added TFA (60 μL). The reaction mixture was stirred for 2 h at room temperature. After azeotropically removal of the solvent with toluene (1 mL × 3), the residue was dissolved in dry DMF (200 μL) and Cy5-NHS in dry DMF (100 μL), $Et_3N$ (10 μL) were added. The reaction mixture was stirred for O/N at room temperature and purified by RP-HPLC (column; Cosmosil 5C18AR2, 250 x 10 mm, mobile phase; $CH_3CN$ : 10 mM TEAA aq. = 50 : 50 → 70 : 30 (linear gradient over 20 min), flow rate; 9 mL/min, detection; UV (220 nm)) to give **MeTz-Asp-Cy5** (1.2 mg, 1.5 μmol) as a blue solid. HR ESI-MS (calc. for $C_{45}H_{51}N_8O_5S$): $[M-H]^-$ = 815.3702 (calc. = 815.3709).

**Synthesis of MeTz-peptides**

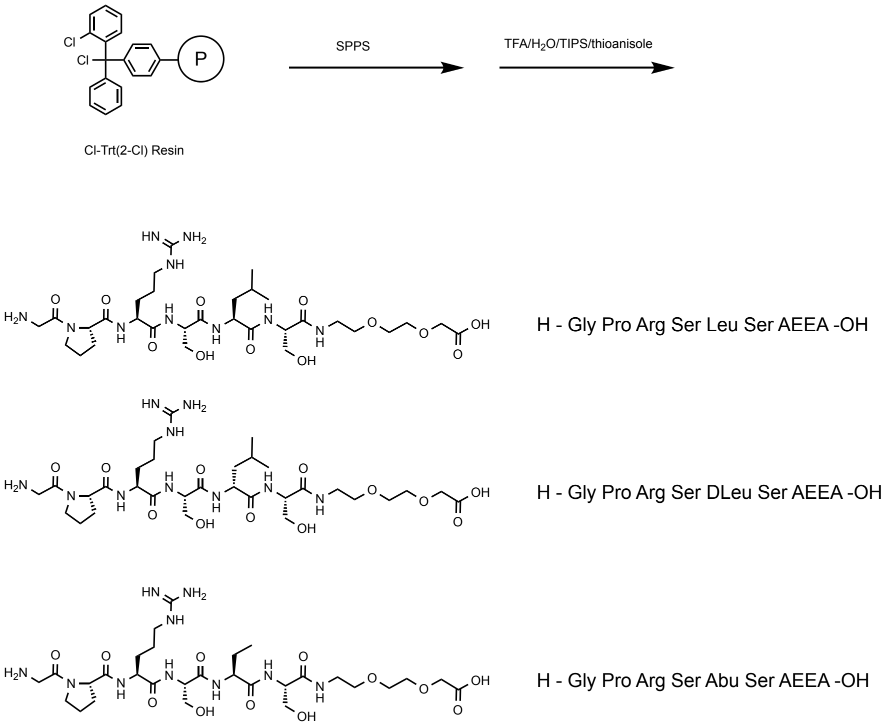


**H-Gly Pro Arg Ser Leu Ser AEEA-OH, H-Gly Pro Arg Ser DLeu Ser AEEA-OH and H-Gly Pro Arg Ser Abu Ser AEEA-OH.** The peptides were synthesized by the stepwise elongation of Fmoc-amino acids on 2-Cl-Trt(2-Cl) resin (0.2 mmol/g, 200 mg, Watanabe chemical co.) according to a reported procedure with Fmoc-AA-OH (5 equiv.) [Fmoc-(2)Abu-OH, Fmoc-AEEA-OH, Fmoc-Arg(Pbf)-OH, Fmoc-Gly-OH, Fmoc-Leu-OH, Fmoc-DLeu-OH, Fmoc-Pro-OH, Fmoc-Ser($Bu^t$)-OH: Pbf, 2,2,4,6-pentamethyldihydrobenzofurane-5-sulfonyl; Trt, trityl; $Bu^t$, *tert*-butyl] using HBTU (5 equiv.), HOBt · $H_2O$ (5 equiv.) and DIPEA (10 equiv.) as coupling reagents in NMP. The protecting groups and the resin were removed by stirring the dried resin for 2 h at room temperature in TFA/$H_2O$/TIPS/thioanisole (85/5/5/5). After removing the resin by filtration, the crude peptides were solidified by adding ice-colded $Et_2O$ and dried under reduced pressure. The crude peptides were purified by RP-HPLC (YMC-pack ODS-A, 250 x 2500 mm, mobile phase; 0.1% TFA/$CH_3CN$ : 0.1% TFA/$H_2O$ = 0:100 → 40:60 (linear gradient over 40 min), flow rate; 10 mL/min, detection; UV absorbance at 220 nm). Yield: **H-Gly Pro Arg Ser Leu Ser AEEA-OH** 33 mg, 0.033 mmol, 83%, **H-Gly Pro Arg Ser DLeu Ser AEEA-OH** 32 mg, 0.032 mmol, 82%, **H-Gly Pro Arg Ser Abu Ser AEEA-OH** 28 mg, 0.029 mmol, 73%. HR-ESI-MS: **H-Gly Pro Arg Ser Leu Ser AEEA-OH** (calc. for $C_{31}H_{56}N_{10}O_{12}$): $[M+H]^+$ = 761.4158 (calc. = 761.4152), **H-Gly Pro Arg Ser DLeu Ser AEEA-OH** (calc. for $C_{31}H_{56}N_{10}O_{12}$): $[M+H]^+$ = 761.4158 (calc. = 761.4152), **H-Gly Pro Arg Ser Abu Ser AEEA-OH** (calc. for $C_{29}H_{52}N_{10}O_{12}$): $[M+H]^+$ = 733.3833 (calc. = 733.3839).

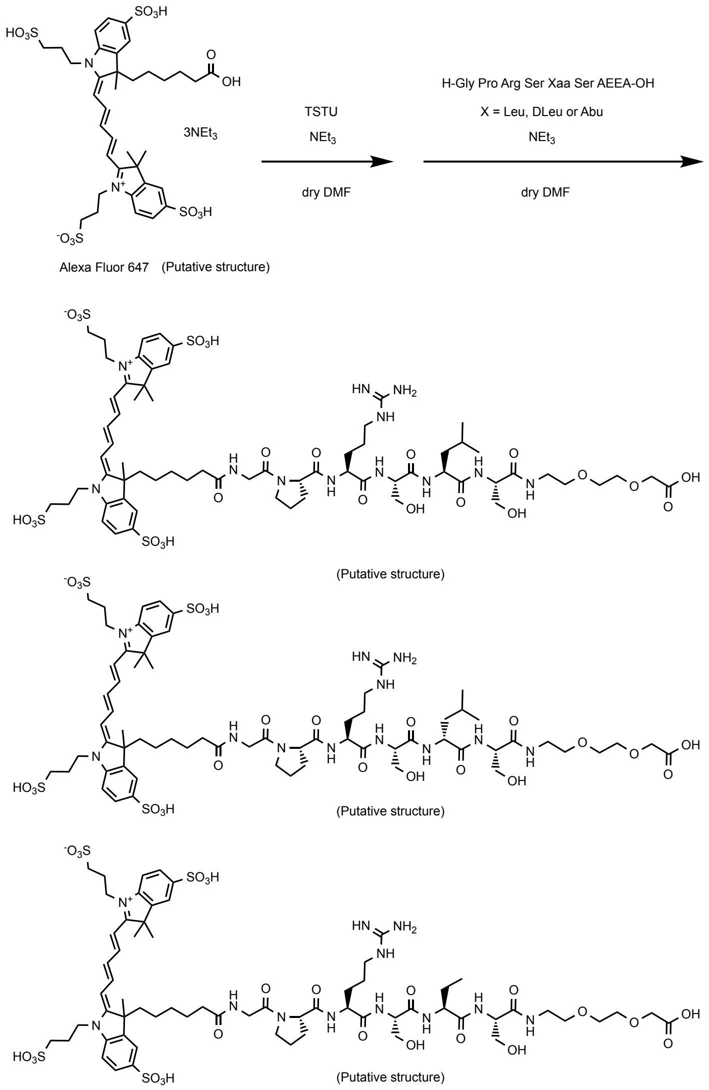


Ax647- Gly Pro Arg Ser Leu Ser AEEA -OH

Ax647- Gly Pro Arg Ser DLeu Ser AEEA -OH

Ax647- Gly Pro Arg Ser Abu Ser AEEA -OH

**Ax647-Gly Pro Arg Ser Leu Ser AEEA-OH, Ax647-Gly Pro Arg Ser DLeu Ser AEEA-OH and Ax647-Gly Pro Arg Ser Abu Ser AEEA-OH.** To a solution of Alexa Fluor 647 (5 mg, 4.3 μmol) in dry DMF (0.2 mL), TSTU (1.9 mg, 6.5 μmol, 1.5 equiv.) and TEA (3.1 μL, 22 μmol, 5 equiv.) dissolved in dry DMF (0.3 mL) were added. The reaction mixture was stirred at room temperature under Ar atmosphere for 2 h. The reaction mixture and TEA (200 μmol ) were added to the peptide (10 μmol) and stirred at room temperature under Ar atmosphere overnight. After removal of DMF by evaporation, the residue was dissolved in 10 mM TEA AcOH buffer (pH 7.0) and purified by RP-HPLC (YMC Triart C18, 10 x 250 mm, mobile phase; $CH_3CN$ : 10 mM TEA AcOH (pH 7.0) = 0:100 → 50:50 (linear gradient over 50 min), flow rate; 3 mL/min, detection; UV absorbance at 220 and 640 nm). Yield (determined by measurement of UV absorbance): **Ax647-Gly Pro Arg Ser Leu Ser AEEA-OH** 2.8 μmol, 64%, **Ax647-Gly Pro Arg Ser DLeu Ser AEEA-OH** 2.9 μmol, 68%, **Ax647-Gly Pro Arg Ser Abu Ser AEEA-OH** 3.3 μmol, 78%. HR-ESI-MS: **Ax647-Gly Pro Arg Ser Leu Ser AEEA-OH** (calc. for $C_{67}H_{100}N_{12}O_{25}S_4$): $[M-H]^-$ = 1599.5735 (calc. = 1599.5733), **Ax647-Gly Pro Arg Ser DLeu Ser AEEA-OH** (calc. for $C_{67}H_{100}N_{12}O_{25}S_4$): $[M-H]^-$ = 1599.5738 (calc. = 1599.5733), **Ax647-Gly Pro Arg Ser Abu Ser AEEA-OH** (calc. for $C_{65}H_{96}N_{12}O_{25}S_4$): $[M-2H]^{2-}$ = 785.2677 (calc. = 785.2673).

Ax647- Gly Pro Arg Ser Xaa Ser AEEA -OH

Xaa = Leu, DLeu or Abu

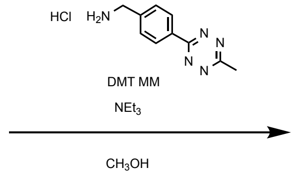


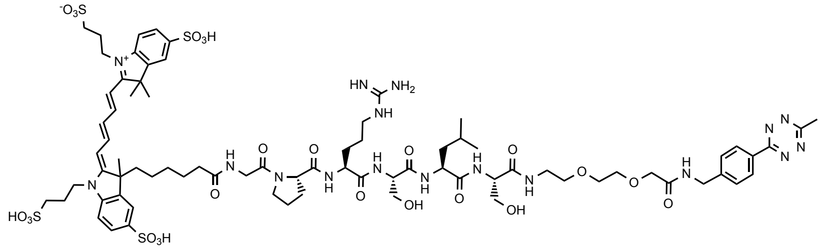

(Putative structure)

Ax647- Gly Pro Arg Ser Leu Ser AEEA -NH-MeTz

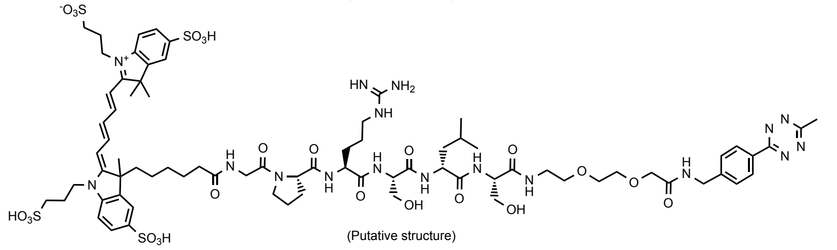

(Putative structure)

Ax647- Gly Pro Arg Ser DLeu Ser AEEA -NH-MeTz

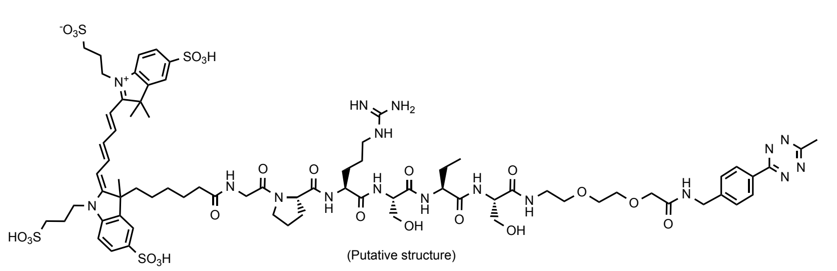

(Putative structure)

Ax647- Gly Pro Arg Ser Abu Ser AEEA -NH-MeTz

**Ax647-Gly Pro Arg Ser Leu Ser AEEA-NH-MeTz (MMP9-Leu), Ax647-Gly Pro Arg Ser DLeu Ser AEEA-NH-MeTz (MMP9-DLeu) and Ax647-Gly Pro Arg Ser Abu Ser AEEA-NH-MeTz (MMP9-Abu).** To a solution of Alexa Fluor 647 conjugated peptide (1.4 µmol) in $CH_3OH$ (1 mL), DMT MM (3.9 mg, 14 µmol, 10 equiv.), methyltetrazine amine HCl (1.7 mg, 7 µmol, 5 equiv.) and TEA (1.0 µL, 7 µmol, 5 equiv.) were added. The reaction mixture was stirred at room temperature overnight. After removal of $CH_3OH$ by evaporation, the residue was dissolved in 10 mM TEA AcOH buffer (pH 7.0) and purified by RP-HPLC (YMC Triart C18, 10 x 250 mm, mobile phase; $CH_3CN$ : 10 mM TEA AcOH (pH 7.0) = 0:100 → 50:50 (linear gradient over 50 min), flow rate; 3 mL/min, detection; UV absorbance at 220 and 640 nm). Yield (determined by measurement of UV absorbance): **Ax647-Gly Pro Arg Ser Leu Ser AEEA-NH-MeTz** 0.92 µmol, 66%, **Ax647-Gly Pro Arg Ser DLeu Ser AEEA-NH-MeTz** 0.95 µmol, 68%, **Ax647-Gly Pro Arg Ser Abu Ser AEEA-NH-MeTz** 0.98 µmol, 70%. HR-ESI-MS: **Ax647-Gly Pro Arg Ser Leu Ser AEEA-NH-MeTz** (calc. for $C_{77}H_{109}N_{17}O_{24}S_4$): $[M-H]^-$ = 1782.6683 (calc. = 1782.6641), **Ax647-Gly Pro Arg Ser DLeu Ser AEEA-NH-MeTz** (calc. for $C_{67}H_{100}N_{12}O_{25}S$): $[M-H]^-$ = 1782.6669 (calc. = 1782.6641), **Ax647-Gly Pro Arg Ser Abu Ser AEEA-OH** (calc. for $C_{75}H_{105}N_{17}O_{24}S_4$): $[M-2H]^{2-}$ = 876.8129 (calc. = 876.8128).

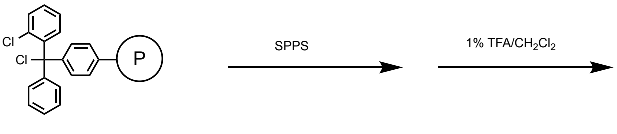


Cl-Trt(2-Cl) Resin

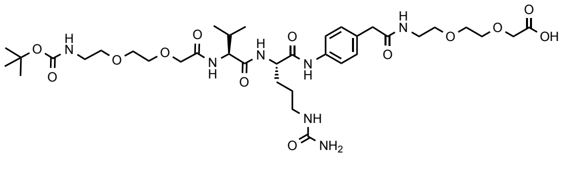

Boc- AEEA Val Cit ABA AEEA -OH

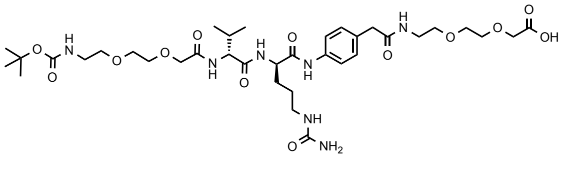

Boc-AEEA DVal DCit ABA AEEA -OH

**Boc-AEEA Val Cit ABA AEEA-OH and Boc-AEEA DVal DCit ABA AEEA-NH-OH.** Fmoc-4-aminobenzeneacetic acid (Fmoc-ABA-OH) was synthesized according to a literature.[S7] The peptides were synthesized by the stepwise elongation of Fmoc-amino acids on 2-Cl-Trt(2-Cl) resin (0.2 mmol/g, 200 mg, Watanabe chemical co.) according to a reported procedure with Fmoc-AA-OH (5 equiv.) [Fmoc-ABA-OH, Fmoc-AEEA-OH, Fmoc-Cit-OH, Fmoc-DCit-OH, Fmoc-Val-OH, Fmoc-DVal-OH using HATU (5 equiv.), HOAt (5 equiv.) and DIPEA (10 equiv.) as coupling reagents in NMP. The peptide was cleaved from the resin by stirring the dried resin for 1 h at room temperature in 1% TFA/$CH_2Cl_2$. After removal of the resin by filtration, the solvent was azeotropically removed with toluene (×3). The crude peptides were dissolved in 10 mM TEA AcOH buffer (pH 7.0) and purified by RP-HPLC (YMC-pack ODS-A, 250 x 2500 mm, mobile phase; $CH_3CN$ : 10 mM TEA AcOH (pH 7.0) = 10:90 → 50:50 (linear gradient over 40 min), flow rate; 10 mL/min, detection; UV absorbance at 220 nm). Yield: **Boc-AEEA Val Cit ABA AEEA-OH** 31 mg, 0.039 mmol, 97%, **Boc-AEEA DVal DCit ABA AEEA-OH** 30 mg, 0.037 mmol, 93%. HR-ESI-MS: **Boc-AEEA Val Cit ABA AEEA-OH** (calc. for $C_{36}H_{59}N_7O_{13}$): $[M-H]^-$ = 796.4106 (calc. = 796.4098), **Boc-AEEA DVal DCit ABA AEEA-OH** (calc. for $C_{36}H_{59}N_7O_{13}$): $[M-H]^-$ = 796.4107 (calc. = 796.4098).

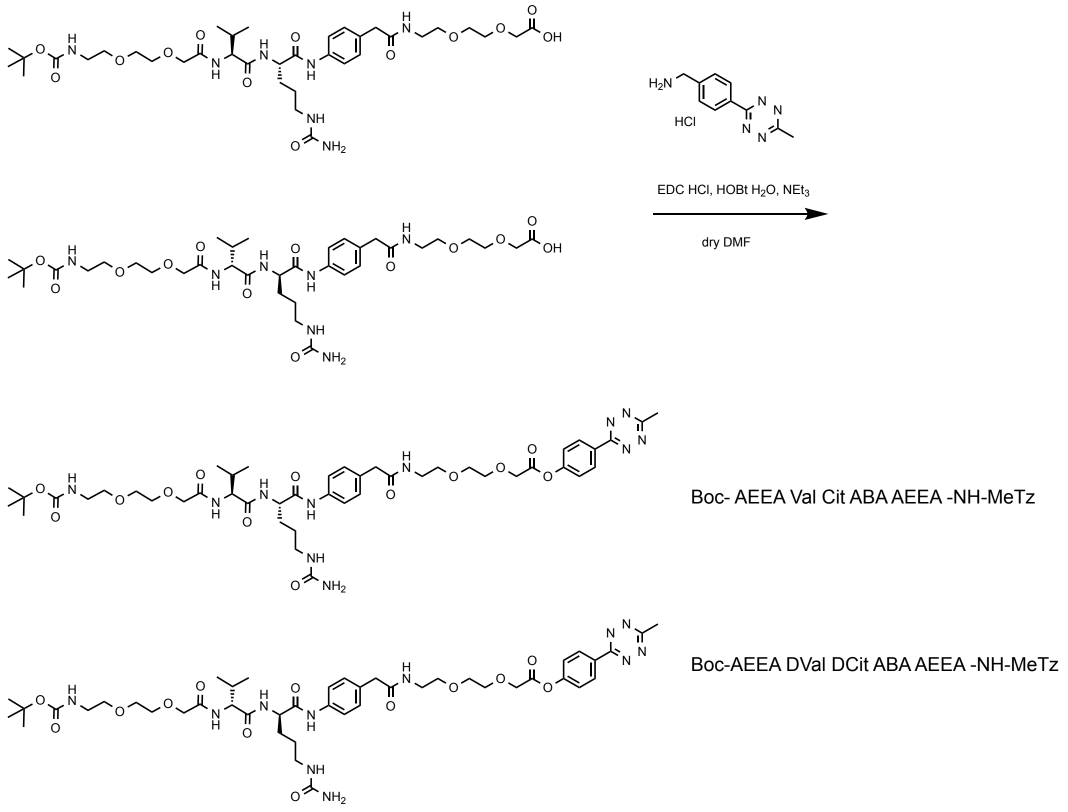


**Boc-AEEA Val Cit ABA AEEA-NH-MeTz and Boc-AEEA DVal DCit ABA AEEA-NH-MeTz.** To a solution of Boc-protected peptide (25 mg, 27.8 μmol, 1.3 equiv.), methyltetrazine amine HCl (5 mg, 21 μmol, 1 equiv.) , EDC HCl (5.3 mg, 27.8 μmol, 1.3 equiv.), HOBt $H_2O$ (4.3 mg, 27.8 μmol, 1.3 equiv.) and TEA (15.5 μL, 111 μmol, 4 equiv.) were added. The reaction mixture was stirred at room temperature overnight. After removal of DMF by evaporation, the residue was dissolved in 10 mM TEA AcOH buffer (pH 7.0) and purified by RP-HPLC (YMC Triart C18, 10 x 250 mm, mobile phase; $CH_3CN$ : 10 mM TEA AcOH (pH 7.0) = 20:80 → 60:40 (linear gradient over 40 min), flow rate; 3 mL/min, detection; UV absorbance at 220 and 640 nm). Yield: **Boc-AEEA Val Cit ABA AEEA-NH-MeTz** 11.0 mg, 11.2 mmol, 53%, **Boc-AEEA DVal DCit ABA AEEA-NH-MeTz** 17.3 mg, 17.6 mmol, 71%. HR-ESI-MS: **Boc-AEEA Val Cit ABA AEEA-NH-MeTz** (calc. for $C_{46}H_{68}N_{12}O_{12}$): $[M-H]^-$ = 979.5008 (calc. = 979.5007), **Boc-AEEA DVal DCit ABA AEEA-NH-MeTz** (calc. for $C_{46}H_{68}N_{12}O_{12}$): $[M-H]^-$ = 979.5006 (calc. = 979.5007).

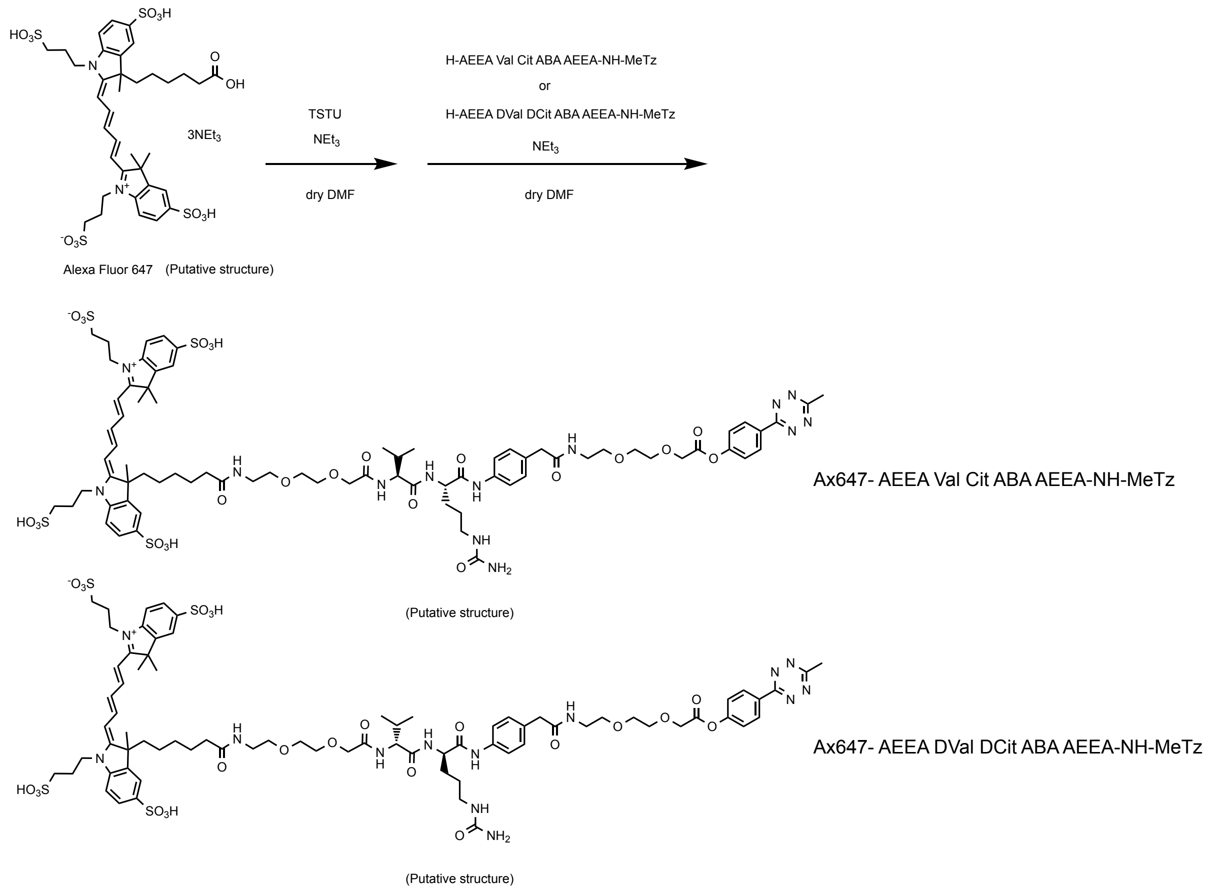


**Ax647-AEEA Val Cit ABA AEEA-NH-MeTz (CatB(L)) and Ax647-AEEA DVal DCit ABA AEEA-NH-MeTz (CatB(D)).** To a solution of methyltetrazine conjugated peptide (2.3 mg, 2.3 μmol) in dry $CH_2Cl_2$ (0.75 mL), TFA (0.25 mL) was added and the reaction mixture was stirred at room temperature for 1 h. After azeotropically removal of the solvent with toluene (×3), the Boc-deprotected peptide was dried under reduced pressure. To a solution of Alexa Fluor 647 (2.5 mg, 2.15 μmol) in dry DMF (0.2 mL), TSTU (0.95 mg, 3.25 μmol, 1.5 equiv.) and TEA (1.6 μL, 11 μmol, 5 equiv.) dissolved in dry DMF (0.3 mL) were added. The reaction mixture was stirred at room temperature under Ar atmosphere for 2 h. The reaction mixture of Alexa Fluor 647 and TEA (9.7 μmol, 61 μmol, 20 eq.) were added to the dried Boc-deprotected peptide and stirred at room temperature under Ar atmosphere overnight. After removal of DMF by evaporation, the residue was dissolved in 10 mM TEA AcOH buffer (pH 7.0) and purified by RP-HPLC (YMC Triart C18, 10 x 250 mm, mobile phase; $CH_3CN$ : 10 mM TEA AcOH (pH 7.0) = 0:100 → 50:50 (linear gradient over 50 min), flow rate; 3 mL/min, detection; UV absorbance at 220 and 640 nm). Yield (determined by measurement of UV absorbance): **Ax647-AEEA Val Cit ABA AEEA-NH-MeTz** 1.26 μmol, 58%, **Ax647-AEEA DVal DCit ABA AEEA-NH-MeTz** 0.84 μmol, 39%. HR-ESI-MS: **Ax647-AEEA Val Cit ABA AEEA-NH-MeTz** (calc. for $C_{77}H_{104}N_{14}O_{23}S_4$): [M-

2H]$^{2-}$ = 859.3069 (calc. = 859.3068), **Ax647-AEEA DVal DCit ABA AEEA-NH-MeTz** (calc. for $C_{77}H_{104}N_{14}O_{23}S_4$): [M-2H]$^{2-}$ = 859.3065 (calc. = 859.3068).
